# Naïve CD8 T cell IFNγ responses to a vacuolar antigen are regulated by an inflammasome-independent NLRP3 pathway and *Toxoplasma gondii* ROP5

**DOI:** 10.1101/2020.01.20.912568

**Authors:** Angel K. Kongsomboonvech, Felipe Rodriguez, Anh L. Diep, Brandon M. Justice, Brayan E. Castallanos, Ana Camejo, Debanjan Mukhopadhyay, Gregory A. Taylor, Masahiro Yamamoto, Jeroen P.J. Saeij, Michael L. Reese, Kirk D.C. Jensen

**Affiliations:** Department of Molecular and Cell Biology, University of California, Merced, Merced, California 95343, USA; Department of Biology, Massachusetts Institute of Technology, Cambridge, Massachusetts 02139, USA; Department of Pathology, Microbiology and Immunology, School of Veterinary Medicine, University of California, Davis, Davis, California 95616, USA; Departments of Medicine; Molecular Genetics and Microbiology; and Immunology; and Center for the Study of Aging and Human Development, Duke University Medical Center, Durham, NC 27710; and Geriatric Research, Education, and Clinical Center, Durham VA Health Care System, Durham, NC 27705, USA; Department of Immunoparasitology, Research Institute for Microbial Diseases, Osaka University, Osaka 565-0871, Japan; Department of Pharmacology, University of Texas, Southwestern Medical Center, Dallas, Texas 75390, USA

**Author notes:** Correspondence: Kirk D.C. Jensen.

## Abstract

Host resistance to *Toxoplasma gondii* relies on CD8 T cell IFNγ responses, which if modulated by the host or parasite could influence chronic infection and parasite transmission between hosts. Since host-parasite interactions that govern this response are not fully elucidated, we investigated requirements for eliciting naïve CD8 T cell IFNγ responses to a vacuolar resident antigen of *T. gondii*, TGD057. Naïve TGD057 antigen-specific CD8 T cells (T57) were isolated from transnuclear mice and responded to parasite-infected bone marrow-derived macrophages (BMDMs) in an antigen-dependent manner, first by producing IL-2 and then IFNγ. T57 IFNγ responses to TGD057 were independent of the parasite’s protein export machinery ASP5 and MYR1. Instead, host immunity pathways downstream of the regulatory Immunity-Related GTPases (IRG), including partial dependence on Guanylate-Binding Proteins, are required. Multiple *T. gondii* ROP5 isoforms and allele types, including ‘avirulent’ ROP5A from clade A and D parasite strains, were able to suppress CD8 T cell IFNγ responses to parasite-infected BMDMs. Phenotypic variance between clades B, C, D, F, and A strains suggest T57 IFNγ differentiation occurs independently of parasite virulence or any known IRG-ROP5 interaction. Consistent with this, removal of ROP5 is not enough to elicit maximal CD8 T cell IFNγ production to parasite-infected cells. Instead, macrophage expression of the pathogen sensors, NLRP3 and to a large extent NLRP1, were absolute requirements. Other members of the conventional inflammasome cascade are only partially required, as revealed by decreased but not abrogated T57 IFNγ responses to parasite-infected ASC, caspase-1/11, and gasdermin D deficient cells. Moreover, IFNγ production was only partially reduced in the absence of IL-12, IL-18 or IL-1R signaling. In summary, *T. gondii* effectors and host machinery that modulate parasitophorous vacuolar membranes, as well as NLR-dependent but inflammasome-independent pathways, determine the full commitment of CD8 T cells IFNγ responses to a vacuolar antigen.

**AUTHOR SUMMARY:** Parasites are excellent “students” of our immune system as they can deflect, antagonize and confuse the immune response making it difficult to vaccinate against these pathogens. In this report, we analyzed how a widespread parasite of mammals, *Toxoplasma gondii,* manipulates an immune cell needed for immunity to many intracellular pathogens, the CD8 T cell. Host pathways that govern CD8 T cell production of the immune protective cytokine, IFNγ, were also explored. We hypothesized the secreted *Toxoplasma* virulence factor, ROP5, work to inhibit the MHC 1 antigen presentation pathway therefore making it difficult for CD8 T cells to see *T. gondii* antigens sequestered inside a parasitophorous vacuole. However, manipulation through *T. gondii* ROP5 does not fully explain how CD8 T cells commit to making IFNγ in response to infection. Importantly, CD8 T cell IFNγ responses to *T. gondii* require the pathogen sensor NLRP3 to be expressed in the infected cell. Other proteins associated with NLRP3 activation, including members of the conventional inflammasome activation cascade pathway, are only partially involved. Our results identify a novel pathway by which NLRP3 regulates T cell function and underscore the need for inflammasome-activating adjuvants in vaccines aimed at inducing CD8 T cell IFNγ responses to parasites.

## INTRODUCTION

*Toxoplasma gondii* is a globally spread intracellular parasite that can infect nearly all warm-blooded vertebrates, including humans. Transmission between hosts occurs following ingestion of oocysts shed from the definitive feline host or predation of chronically infected animals harboring infectious ‘tissue cysts’. Immune modulation by the parasite during the first weeks of infection is therefore critical for *T. gondii* to establish latency and lifecycle progression. The parasite accomplishes this by hiding and manipulating the immune system from within a specialized parasitophorous vacuole (PV) that is created during invasion. *T. gondii* releases ‘effector’ proteins from secretory organelles, including rhoptry proteins (ROP) that are injected into the host cytosol upon invasion, as well as dense granules (GRA) that are secreted into the lumen of the PV and aid its internal structure and formation. Many of these secreted ‘effectors’ manipulate host cell signaling pathways and shield the PV from host immune attack (1). In mice, *T. gondii* uses several ROP and GRA proteins to antagonize the host’s Immunity-Related GTPases (IRGs) which target and compromise the PV (2). ROP5 is encoded by a multi-gene variable family of pseudokinases and can directly bind to and induce allosteric changes in host IRGs (3), presenting them for phosphorylation by the ROP18 (4, 5) and ROP17 kinases (6). The process of phosphorylation inactivates host IRGs, preventing them from assembling on the surface of the PV, which in turn allows the parasite to replicate (7, 8). Genetic variations in ROP5 and ROP18 largely explain parasite strain differences in mouse virulence (9–13), highlighting the importance of the IRG system in the control of *T. gondii* infection. IRGs also regulate the recruitment of Guanylate Binding Proteins (GBPs) and autophagy machinery to the PV membrane (PVM), both of which contribute to cell autonomous immunity to *T. gondii* (14, 15).

Since IRGs and GBPs are induced transcriptionally following stimulation with IFNγ (16), immune cells that produce IFNγ are critically important for resistance to *T. gondii* (17). CD8 T cell IFNγ responses are required for host survival to *T. gondii* infections (18–21) and to prevent reactivation of the dormant form (22, 23). In vaccinated or chronically infected mice, IFNγ and CD8 T cells are primarily responsible for protection against lethal secondary infections (24, 25). However, most *T. gondii* strains that express virulent alleles of ROP5 and ROP18 evade the host’s immunological memory response and superinfect the brains of challenged survivors (26), implicating that sterile immunity to *T. gondii* may be difficult to achieve, as noted for other parasitic pathogens (27). Whether *T. gondii* manipulates induction of the host’s IFNγ response to prolong its survival is unknown, but could represent a general strategy to promote persistence and latency, as noted for numerous viral pathogens in the presence of clonally expanded antigen-specific CD8 T cells (28).

In order for naïve CD8 T cells to become IFNγ producers, they must first be activated by peptides derived from the host’s MHC 1 antigen presentation pathway and then receive cues from the environment or other immune cells to differentiate into IFNγ-producing cells. The question of MHC 1 antigen presentation for *T. gondii* antigens has largely been addressed using two experimental systems. One analyzes antigen-specific CD8 T cell responses to parasite strains expressing the model antigen, chicken ovalbumin (OVA) (29, 30), and the other analyzes responses to *T. gondii* immune-dominant antigen GRA6, encoded by type II strains (31, 32). From these studies, it is appreciated that active cell invasion by *T. gondii*, rather than phagocytosis of invasion-blocked or heat-killed parasites, is required to stimulate host CD8 T cells (30,33,34). The antigen must be in the parasite’s secretory pathway (30, 35), degraded by host cytosolic proteasomes (31, 36), transported via the endoplasmic reticulum (ER) TAP1/2 translocon (29–31,34), and eventually loaded onto MHC 1 molecules. Although dense granules and rhoptry proteins access the host cytosol where MHC 1 antigen processing readily occurs, antigens targeted to the dense granule secretory pathway elicit a greater CD8 T cell response (35). The PV is therefore a suitable platform for MHC 1 antigen presentation, which is remarkable given the PV of *T. gondii* does not initially fuse with host organelles (37, 38), nor is contained within the conventional endocytic compartments of the cell.

The mechanism by which the immune system gains access to PV antigens of *T. gondii* has remained an active area of research, notwithstanding for its implication in vaccine development (39) and the ability of *T. gondii* to elicit anti-tumor responses (40). In the case of *T. gondii* GRA6, it must be integrated in the PVM (41), where its C-terminal epitope (32) is exposed to the host cytosol (42) and degraded by unknown proteases. For the MHC 1 antigen presentation of transgenic OVA expressed in the PV lumen of *T. gondii*, two general mechanisms have been reported. Fusion between the PVM and the host ER (34), or ER-derived Golgi Intermediate Compartments (ER-GICs) promotes OVA-specific CD8 T cell activation (43). In this scenario, through a SNARE Sec22b-dependent mechanism, the host’s MHC 1 antigen-processing machinery gains access to the PV whereby it shuttles parasite proteins into the cytosol for antigen processing. In a second mechanism, though not mutually exclusive, the PVM is compromised by the host’s IRGs and selective autophagy systems therefore allowing OVA antigen release (30, 44). Via ROP5 and ROP18, *T. gondii* can bypass IRGs activity and presumably MHC 1 antigen presentation by sequestering OVA inside an intact PV (45). However, several dense granule proteins are also implicated, signifying multiple non-redundant pathways may regulate MHC 1 antigen presentation of PV antigens (45). Whether lessons learned from GRA6 and OVA extend to other antigens or parasite genetic backgrounds is unknown.

In addition to activation by antigen, CD8 T cells need proper co-stimulation (46, 47) and IL-12 signaling to fully commit to IFNγ production during *T. gondii* infection (18,48,49). IL-18 is important for host survival during acute *T. gondii* infection (50), and is released following parasite detection and inflammasome activation by the pathogen sensors NLRP3 and NLRP1 (50, 51). Inflammasome matured IL-18 is important for IFNγ production by CD4 T cells but is apparently dispensable for CD8 T cell IFNγ-production during acute *T. gondii* infection (52). Whether the inflammasome contributes to CD8 T cell activation or differentiation in different contexts or stages of *T. gondii* infection is unclear.

Given the parasite’s need to establish latency and the host’s dependence on CD8 T cells for immunity, we asked whether *T. gondii* has evolved to manipulate CD8 T cell IFNγ responses to an endogenous antigen, and whether certain *T. gondii* genotypes are defined by their ability to induce or repress the production of this immune-protective cytokine. Through the use of T cell receptor transnuclear and IFNγ reporter mice, host and parasite requirements were defined for the induction of IFNγ-producing CD8 T cells to a conserved vacuolar antigen of *T. gondii*, TGD057. Here we report that TGD057-specific CD8 T cell responses are independent of the parasite’s PV-export machinery, and like previous findings with OVA-engineered *T. gondii* strains, the IRG pathway is required. Multiple ROP5 isoforms suppress this response, including ROP5A which lacks a defined function or interaction with host IRGs. An analysis of parasite strains spanning twelve haplogroups suggests IFNγ production is manipulated by *T. gondii* independent of any known IRG-ROP5 interaction or parasite virulence factor. Importantly, an NLRP3-dependent but inflammasome complex-independent pathway is required for inducing maximal CD8 T cell IFNγ responses to *T. gondii* infected cells. Our findings point to novel host-parasite interactions by which IRGs and NLRP3 shape CD8 T cell IFNγ responses to an intracellular pathogen.

## RESULTS

### Naïve CD8 T cells respond to the vacuolar antigen TGD057 with a robust IFNγ response

To determine host and parasite requirements for eliciting antigen-specific CD8 T cell responses to *T. gondii*, we took advantage of ‘T57’ transnuclear mice which were cloned from the nucleus of a single tetramer-positive *T. gondii*-specific CD8 T cell. T57 T cells from these mice have a single T cell receptor (TCR) specificity for the TGD057_96-103_ epitope presented by H-2K^b^ MHC 1 (49, 53), and when adoptively transferred, confer resistance to infection with a type II strain (53). TGD057 is a protein with unknown function but is predicted to be in the parasite’s secretory pathway (54), and when deleted does not negatively impact parasite fitness (not shown) (phenotype score 2.1) (55). Importantly, *TGD057* (Tg_215980) is highly expressed (ToxoDB) and the peptide epitope is conserved between strains (Fig S1), facilitating comparative analyses of naïve CD8 T cell responses to parasite strains which may differ in immune modulation. In our experimental setup (i.e. the T cell activation ‘T57 assay’) (Fig 1A), bone marrow-derived macrophages (BMDMs) are infected with *T. gondii*, co-cultured with splenocytes and lymph node cells from naïve transnuclear T57 mice, and CD8 T cell activation markers or effector cytokines in the supernatant are measured. Reflecting early T cell activation events culminating in calcium-dependent NFAT activation of the IL-2 gene (56), this cytokine is produced as early as 24 hours post addition of T cells to parasite-infected BMDMs (Fig 1B). In contrast and consistent with their naïve state, the T57 IFNγ response develops with time, and is maximally detected at 48 hours (Fig 1C). *T. gondii* infection elicits strong Tc1 responses to TGD057 (49) and other antigens (57), as such IL-17 is only marginally detected in this system (Fig S2). The measured phenotypes are antigen-specific, because the cytokine response is abolished in response to *Δtgd057* strains which do not express the antigen (Fig 1D-1E). Additionally, T57 CD8 T cells fail to upregulate the early activation marker CD69 in response to *Δtgd057* (not shown). Previous observations from sub-cellular fractionation and immuno-florescence studies have identified TGD057 both within the PV of infected cells (41) and to the cytoskeleton region of the parasite (48). Three-dimensional mass spec LOPIT analysis (Location of Organelle Proteins by Isotype Tagging) posits TGD057 to dense granules but lacks a strict assignment to any one organelle, the latter observation being consistent with most cytoskeleton network associated proteins (58). An endotagged RH_tgd057_-HA strain was generated and TGD057 is always found within PVM defined by GRA7 staining vacuoles (Fig 1F), demonstrating TGD057, in its natural state, stays inside the PV. In summary, the T57 system allows analysis of naïve CD8 T cell responses to an endogenous vacuolar antigen of *T. gondii*.

**Figure 1:**
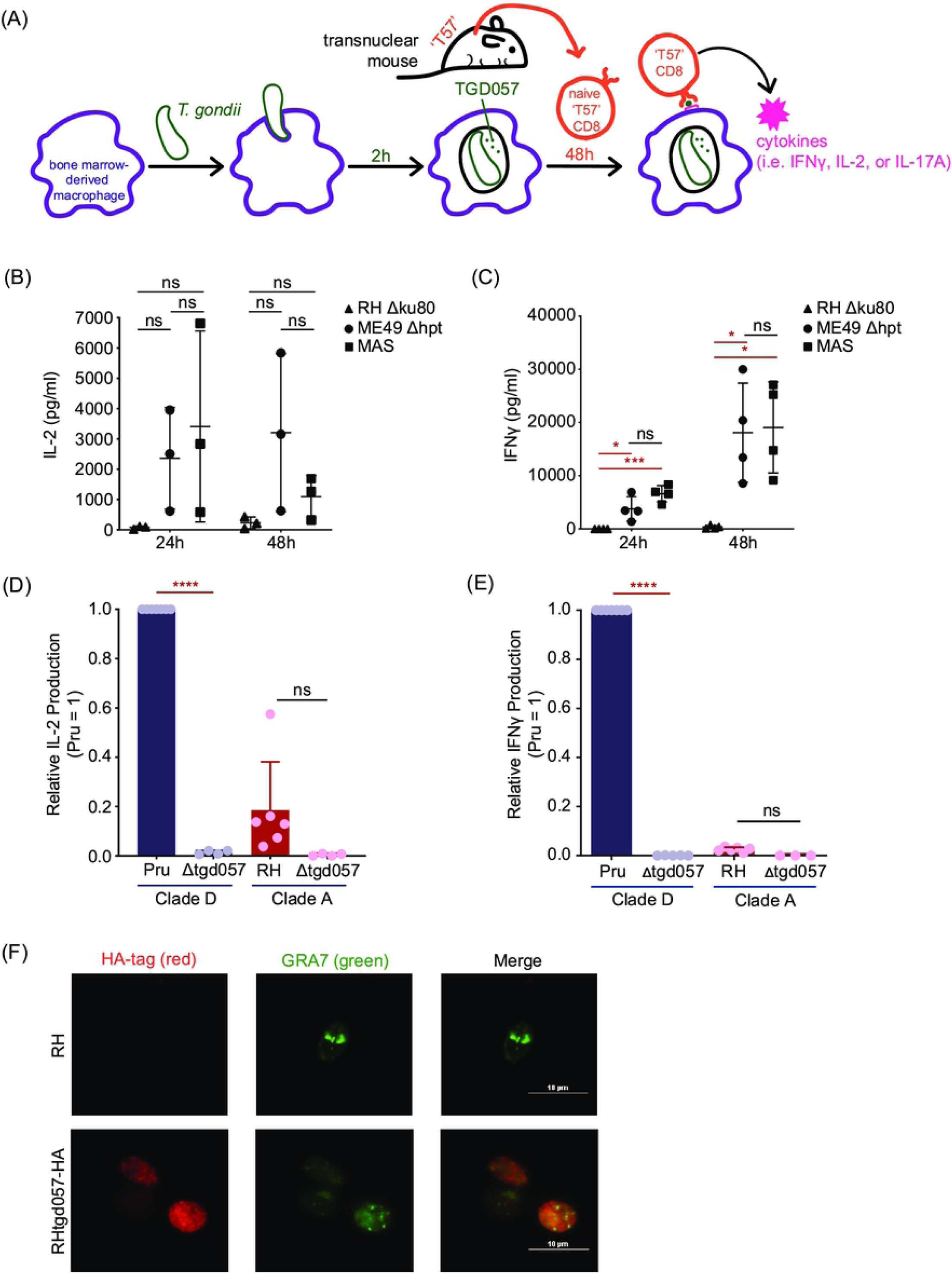
A model to study naïve CD8 T cell responses to a *T. gondii* vacuolar antigen, TGD057. **(A)** Schematic of the ‘T57 assay’. Bone marrow-derived macrophages (BMDMs) are infected with *T. gondii* and 2h later, naïve T57 CD8 T cells obtained from transnuclear mice and added to the infected macrophages. T57 T cells bear antigen receptor specificity for a natural *T. gondii* antigen, the processed TGD057_96-103_ peptide in complex with MHC 1 K^b^. Supernatant from the co-culture is then harvested and cytokine concentrations are analyzed by ELISA. **(B-C)** TGD057-specific CD8 T cell responses to *T. gondii*-infected macrophages were measured over time to the indicated parasite strains. At 24h and 48h time points, IFNγ and IL-2 was measured by ELISA. Average of 3-4 experiments + SD (standard deviation) are plotted; each dot represents the result from an individual experiment. Statistical analysis comparing parasite strain-differences were performed by two-way ANOVA with Bonferroni’s correction; * p ≤ 0.05, *** p ≤ 0.001, and ns non-significant. **(D-E)** Parental and *Δtgd057 T. gondii* strains were assayed for the CD8 T cell response as described in Fig 1A. T57 IFNγ and IL-2 responses at 48h, analyzed by ELISA, are normalized to that of the clade D wildtype strain (Pru). Average of 3-5 experiments + SD are shown, each dot represents the results from an individual experiment. Statistical analysis between parental and knockout strains is performed by an unpaired t-test; **** p ≤ 0.0001, ns non-significant. **(F)** Human foreskin fibroblasts (HFFs) were infected with RH or an RH_tgd057_-HA endotagged strain. After 20 hours of infection, the samples were fixed, permeabilized and the tagged TGD057-HA was visualized in rat anti-HA antibodies, visualized in red. PVM is indicated by presence of the PVM integral and PV luminal dense granule protein, GRA7, visualized in green. A representative immunofluorescence image is shown.

### TGD057 antigen acquisition is not dependent on the parasite’s protein export pathway

To understand how *T. gondii* vacuolar antigens might escape from the vacuole and enter the host’s MHC 1 antigen presentation pathway, the parasite’s export machinery was explored. One way for vacuolar proteins to enter the host cytosol is through parasite-mediated export across the PV membrane (PVM). Dense granule proteins that reside within the PV can leave the vacuole through *T. gondii*’s export machinery, which includes the Golgi-resident protein aspartyl protease, ASP5 (59, 60), and the PVM-integrated translocon protein MYR1 (61). *T. gondii* ASP5 is an orthologue of *Plasmodium* protease Plasmepsin V which recognizes a *<u>P</u>lasmodium* <u>ex</u>port <u>el</u>ement (PEXEL) motif (RxLxE/Q/D) (62) and cleaves after the leucine (RxL↓xE/Q/D) preparing PEXEL-bearing proteins for export across the PVM into the host erythrocyte (63). Like Plasmepsin V, *T. gondii* ASP5 recognizes and cleaves a PEXEL-like motif (RRL↓XX) (*<u>T</u>oxoplasma* <u>ex</u>port <u>el</u>ement or ‘TEXEL’ motif) (60), its protease function is necessary for the export of all known exported PV proteins (64). For example, GRA16 contains an RRL↓XX sequence, is cleaved by ASP5, and utilizes the MYR1 translocon complex for protein export (60, 65). As TGD057 contains an RRL↓XX sequence, we hypothesized ASP5 and/or MYR1 may be involved in the export of TGD057 from the PV, leading to MHC 1 antigen presentation and T57 antigen-specific CD8 T cell responses. To test this, the TGD057-specific CD8 T cell response to *Δasp5* and *Δmyr1* strains was measured as previously described in Figure 1. In contrast to the hypothesis, ME49 *Δasp5* induced a higher CD8 T cell response compared to that of wildtype ME49 (Fig 2). Moreover, T57 responses to ME49, ME49 *Δmyr1* and *MYR1* complementation strains (ME49 *Δmyr1::MYR1*) were comparable (Fig 2). Since the T57 cytokine response to the type I RH strain was uniformly low (Fig 1B-1C), inferring requirements for export machinery using these parasite strains was uninformative. Nonetheless, CD8 T cell responses to the type II strain do not require ASP5 and MYR1, suggesting protein export from the PV is not necessary for MHC 1 antigen presentation of the vacuolar TGD057 antigen.

**Figure 2:**
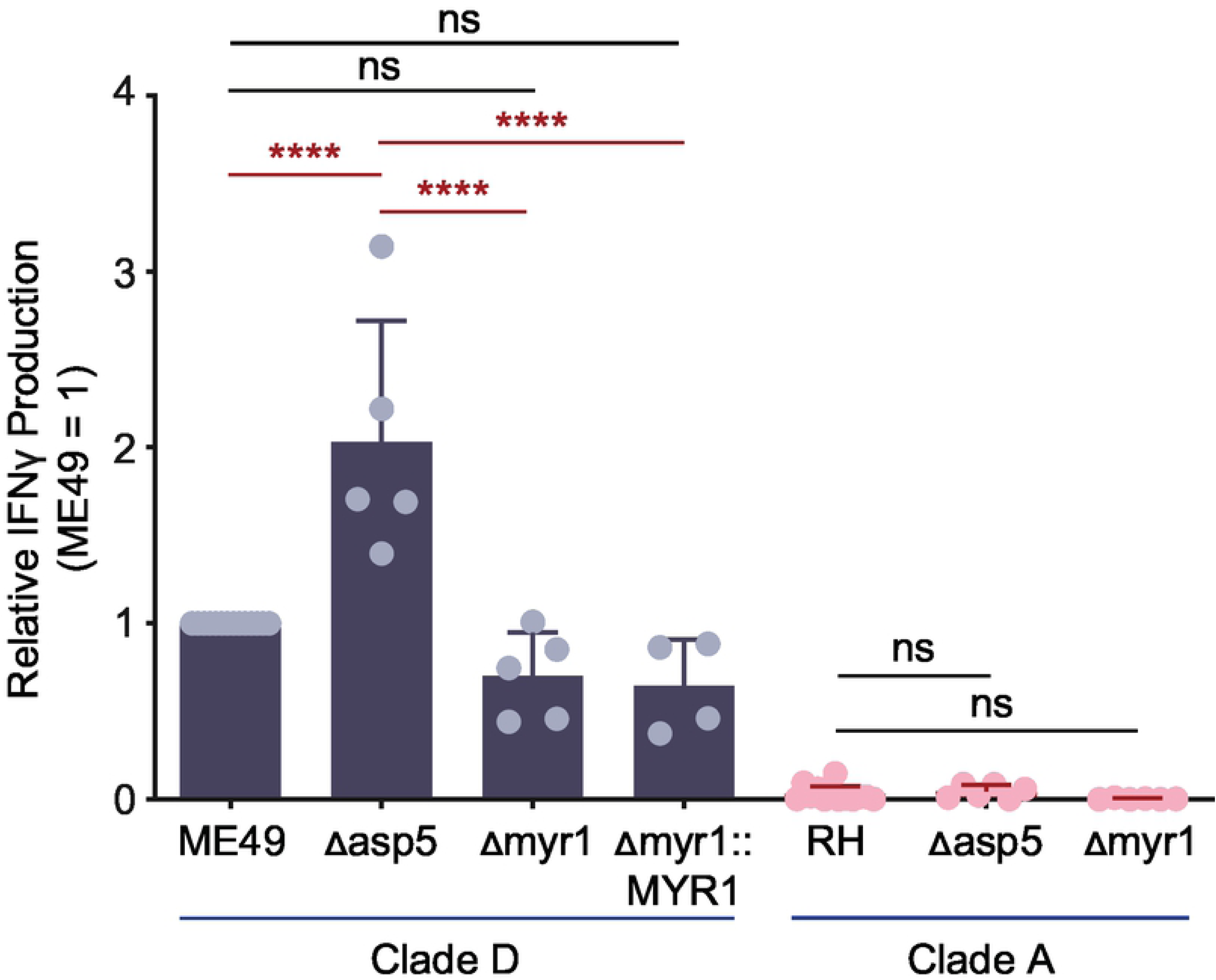
TGD057-speicifc CD8 T cell IFNγ responses do not require the parasite’s export machinery. The *Δasp5* and *Δmyr1 T. gondii* strains listed were assayed for host CD8 T cell response as previously described in Fig 1A. The IFNγ response at 48h, as analyzed by ELISA, is normalized to that of the clade D wildtype strain (ME49). Average of 4-6 experiments + SD are shown, each dot represents the results from one experiment. Statistical analysis was performed using one-way ANOVA with Bonferroni’s correction comparing to parental strains; **** p ≤ 0.0001, ns non-significant.

### CD8 T cell IFNγ responses to TGD057 require host machinery downstream of the regulatory IRGs

Instead we hypothesized the T57 CD8 T cell response requires PV disruption and is IRG-mediated, as implicated from studies of OT1 CD8 T cell responses, or hybridoma derivatives, to parasite strains that express the model OVA antigen in the PV lumen (45,66,67). To this end, T57 assays were performed with various BMDMs that are defective in IRG function. In the experimental setup, IFNγ is derived from activated T57 cells and is predicted to induce IRG expression in WT but not *Stat1-/-* or *Ifngr-/-* macrophages. Consistent with this supposition, the TGD057-specific CD8 T cell response to the ME49 strain was nearly abolished in the absence of IFNγ-STAT1 signaling (Fig 3A). In mice, there are 23 IRGs that can be separated into two subfamilies: 1) the effector IRGs (or ‘GKS class’ based on an amino acid motif in their GTP binding P-loop) and 2) the regulatory IRGs (‘GMS class’) (68). Whereas effector IRGs bind the vacuolar membrane of the PV (69, 70) and mediate membrane destruction via GTP hydrolysis (71), regulatory IRGs (or ‘IRGMs’) localize to host cellular organelles preventing effector IRGs from destroying host membranes (72, 73). Regulatory IRGMs bind to effector IRGs keeping them in their GDP bound inactive state (74), in a manner similar to *T. gondii* ROP5 (3,4,75). In the absence of IRGMs, effector IRGs fail to localize to pathogen PVs (74, 76) and pathogen restriction is lost (77). In mice there are three regulatory IRGs (IRGM-1, −2 and −3) and *irgm1-/-* or *irgm1/3-/-* double knockout macrophages were analyzed. Similar to the OVA system, the TGD057-specific CD8 T cell response to stimulatory parasite strains, such as type II ME49 and the atypical strain MAS, require the activity of regulatory IRGMs (Fig 3A-3B). In addition, IFNγ-inducible Guanylate-Binding Proteins (GBPs) localize to the PV in an IRGM-dependent manner (72) and mediate *T. gondii* resistance (15). GBPs encoded on murine chromosome 3 (GBP^chr3^) are involved in PV disruption, promote effector IRG recruitment to the PV of *T. gondii* (67, 78), and once compromised will attack the parasite’s plasma membrane, decreasing its fitness (14). To test whether GBPs are required for TGD057-specific CD8 T cell responses, GBP^chr3^-deficient BMDMs lacking *Gbp1*, *Gbp2*, *Gbp3*, *Gbp5*, and *Gbp7* were screened (67). GBP^chr3^ were not significantly involved in promoting TGD057-specific CD8 T cell IFNγ responses to the ME49 strain (Fig 3C), but were partially required for the response to the atypical strain MAS (Fig 3D). Altogether, these observations are consistent with a model in which the PVM is compromised by host machinery downstream of regulatory IRGs, including GBPs (72) and likely other immunity genes, that mediate vacuolar antigen escape from the PV and entry into the host’s MHC 1 antigen-presentation pathway.

**Figure 3:**
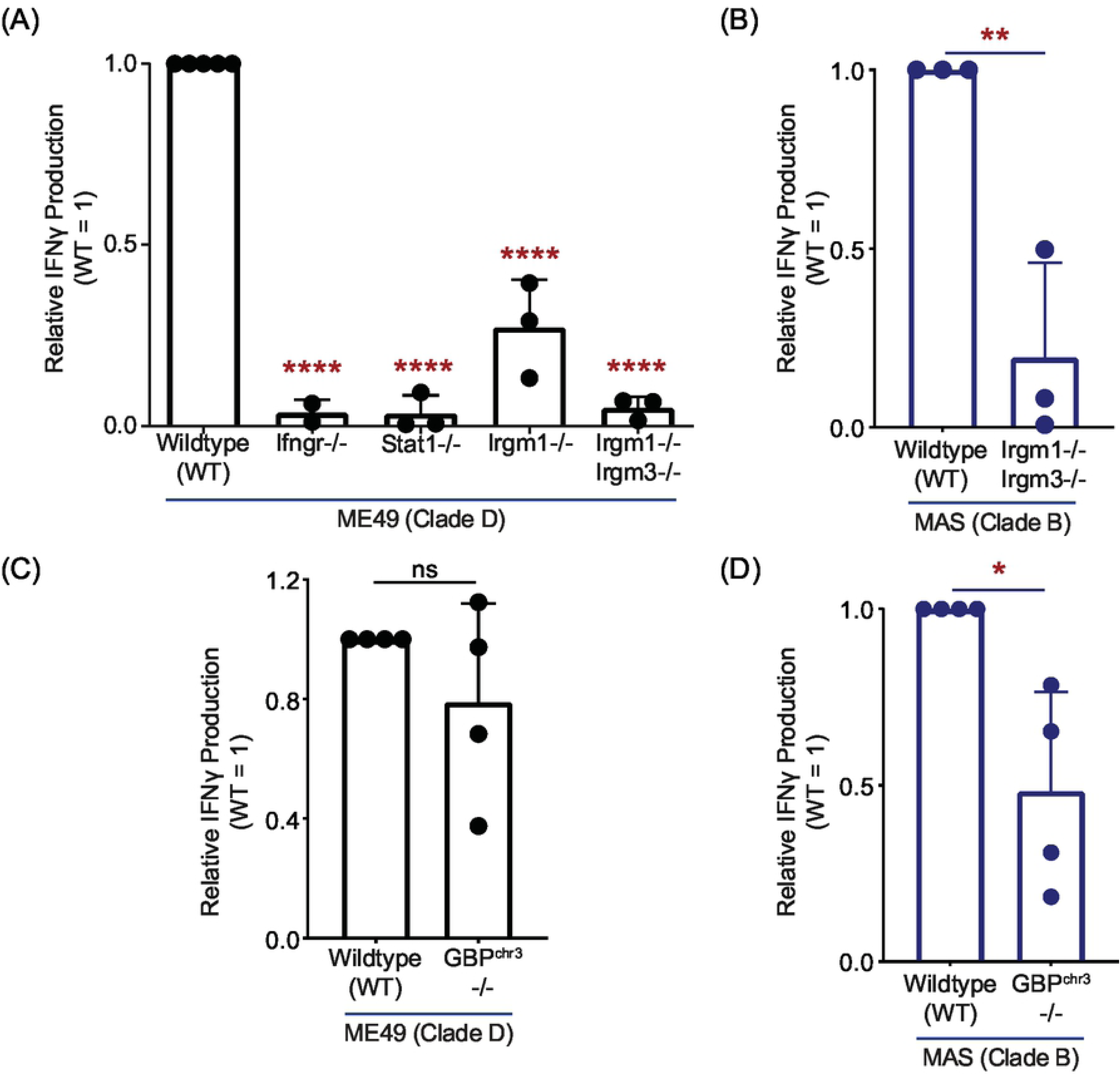
TGD057-specific CD8 T cell IFNγ responses are partially dependent on host GBPs but entirely dependent on regulatory IRGs. **(A-B)** BMDMs with indicated gene deletion (-/-) were infected with the clade D ME49, or **(C-D)** clade B MAS strain. T57 T cell IFNγ responses to TGD057 were analyzed by ELISA at 48h and normalized to the response elicited by infected wildtype (WT) BMDMs. Average of 2-4 experiments + SD are shown, each dot represents the result from an individual experiment. Statistical analyses were performed by one-way ANOVA with Bonferroni’s correction **(A)** or unpaired two-tailed t-tests **(B-D)**; * p ≤ 0.05, ** p ≤ 0.01, **** p ≤ 0.0001, ns non-significant.

### Multiple ROP5 isoforms of *T. gondii* suppress the CD8 T cell response to TGD057

Previous studies have shown that *T. gondii* virulence in mice is determined by parasite effectors that protect the PV from host immune attack. Alleles and isoforms from virulent strains of the rhoptry pseudokinase ROP5 (3,5,9,12,13,79), ROP18 (4,7,10,75,80) and ROP17 kinases (4, 75), are noted for their ability to inhibit the destructive functions of IRGs at the PVM, including Irgb6 and Irga6 (2–4,74,75). These rhoptry proteins also impact the association of GBPs with the PV (67,78,81–83). Thus, we reasoned the host CD8 T cell response to TGD057 would be antagonized by some or all of these secreted effectors. Indeed, when ROP5 is not expressed in the type I RH strain (clade A, RH *Δrop5*), the CD8 T cell IFNγ and IL-2 response is robust (Fig 4, Fig S3A), and on average, the IFNγ response is half of that elicited by the stimulatory type II strains (Fig 4). The *ROP5* locus consists of *ROP5A*, *ROP5B* and *ROP5C* genes, which differ in copy number between strains, and is under diversifying selection (3,9,12,13,75,79). A series of RH Δ*rop5* strains complemented with one or two copies of *ROP5B* or *ROP5A* isoforms from clade A, or a single *ROP5A* isoform from the type II genetic background (clade D) was analyzed. Importantly, all RH *Δrop5*+*ROP5* complementation strains phenocopied the T57 response to the parental RH strain (Fig 4). Since ROP5A inhibits the T57 response (Fig 4), we infer Irgb6 and Irga6 are likely not responsible for this phenotype because the ROP5A isoform does not inhibit Irgb6 or Irga6 coating of the PVM (4), nor Irga6 oligomerization (3). Consistent with this supposition, ROP5B but not ROP5A_D_ nor ROP5A isoforms inhibit Irgb6-PV association (Fig S4). Of note, the sequestosome-1 (p62) associates with the PV in an IFNγ-induced IRGM-dependent manner to promote OT1 T cell responses to OVA-expressing *T. gondii* strains (44). However, p62-PV association was not inhibited by ROP5A but instead phenocopied the PV-association patterns of Irgb6 (Fig S4). The ROP18 (4) and ROP17 kinases (6) phosphorylate and inactivate Irga6 and Irgb6. Although there were slight increases in the T57 IFNγ response to the RH *Δrop17* and RH *Δrop18* strains, these responses were not significantly different compared to that of the parental RH strain (Fig 4). Altogether, the data show that multiple ROP5 alleles and isoforms can suppress the T57 response to the TGD057 antigen, and this most likely by a mechanism independent of host Irgb6, Irga6 and p62 localization to the PV.

**Figure 4:**
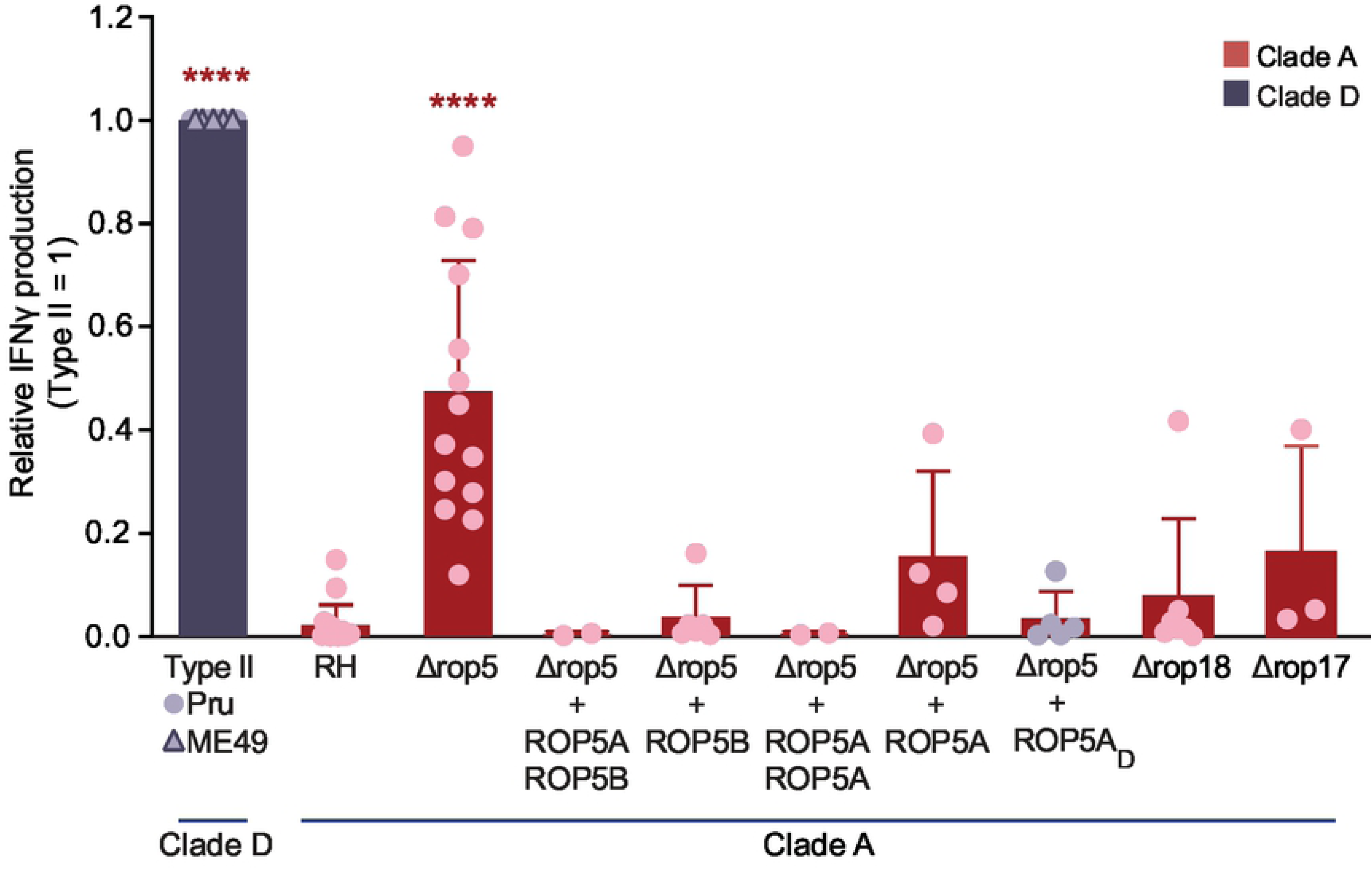
Multiple ROP5 isoforms inhibit TGD057-specific CD8 T cell IFNγ responses to *T. gondii*. T57 CD8 T cell IFNγ responses to the RH and RH *Δrop5* strains, including various ROP5A and/or ROP5B complementation strains from clade A and one from clade D (RH *Δrop5* +*ROP5A_D_*), were analyzed as described in Fig 1A. Additionally, T57 IFNγ responses to RH *Δrop17* and RH *Δrop18* strains were determined. IFNγ was detected by ELISA at 48h and normalized to the response elicited by clade D strains, Pru (O) or ME49 (Δ). Average of 2-14 experiments + SD is shown, each dot represents the result from an individual experiment. Statistical analysis was performed using one-way ANOVA with Bonferroni’s correction comparing all strains to the RH strain, only type II and RH *Δrop5* strains proved significantly different from RH over multiple experiments; **** p ≤ 0.0001.

### ROP5-expressing clade A strains confer low TGD057-specific CD8 T cell IFNγ responses

Due to the conserved nature of the TGD057 peptide epitope (Fig S1), a unique opportunity arose to explore the development of CD8 T cell IFNγ responses to multiple parasite strains spanning the genetic diversity of *T. gondii* (84). Any observed trend between *T. gondii* virulence, genetic background and the T cell response may offer clues to possible parasite immune modulation and adaptation to immune pressure incurred by CD8 T cells. Among the Eurasian clonal strains (types I, II, III) and North American isolates from haplogroups XI (COUGAR) and XII (B73), the virulent type I strains (GT1, RH) induced the lowest CD8 T cell response while intermediate virulent types II (Pru, ME49), XI (COUGAR) and XII (B73) and low virulent type III (CEP) strains, induced relatively high CD8 T cell IFNγ and IL-2 responses (Fig 1B-1C, 5A, S3B). ‘Atypical’ strains, many of which are endemic to South America and highly virulent in laboratory mice (FOU, CAST, MAS, TgCatBr5, P89, GUY-MAT, GUY-DOS, GUY-KOE, RUB, and VAND), differed dramatically in eliciting T57 cytokine responses (Fig 5A, S3B), signifying that parasite virulence is not a sole predictor of CD8 T cell activation. Instead a unique phenotypic pattern emerged, in which clade A strains (Types I, HG VI and VII) conferred low T57 cytokine responses while most other strains from clades B, C, D and F had potential to induce high cytokine responses (Fig 5A). Consistent with previous results, T57 responses did not correlate with known Irgb6 and/or Irga6-PV associations of these strains. For example, a low percentage (∼10%) of Irgb6 recruitment to the PV is observed for the MAS strain, yet a high CD8 T cell IFNγ response is induced (Fig 5A), similar in magnitude to that of type II clade D strains (Figs 1C, 5A) whose PVs are highly decorated with Irgb6 (∼45%) (75). Even among highly virulent type I GT1 and Guyanan strains (i.e. GUY-KOE, GUY-MAT, GUY-DOS, VAND), where approximately 25% or less Irgb6-PV coating is observed (75), the CD8 T cell responses to these strains differ dramatically (Fig 5A, S3B). Furthermore, one notable outlier among clade A strains is the relatively high T57 response to BOF (Fig 5B, S3B). BOF encodes a single copy of *ROP5B* that is marginally expressed (75). When BOF is complemented with the LC37 cosmid, which encodes the entire ROP5 locus from the clade A type I genetic background, the CD8 T cell IFNγ response is largely reduced (Fig 5B). We infer from these assays the clade A genetic background inhibits T57 IFNγ responses, but the identity of the ROP5-host interacting partner and why this genetic background leads to repressed responses is currently unknown.

**Figure 5:**
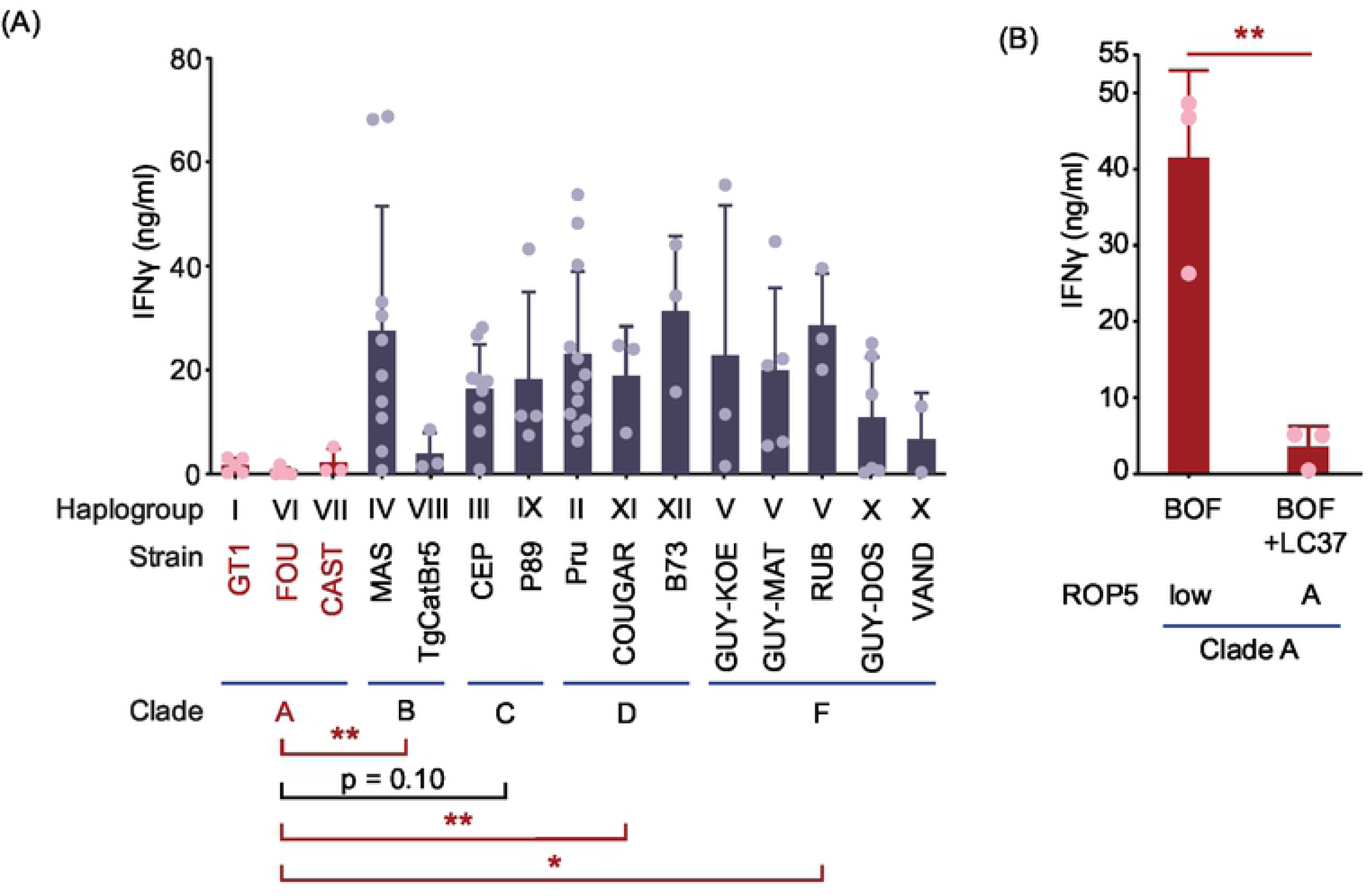
Low CD8 T cell IFNγ responses to ROP5-expressing clade A strains of *T. gondii.* **(A)** The T57 CD8 T cell IFNγ response to BMDMs infected with various *T. gondii* strains from most haplogroups (HG) were analyzed, including the clonal (types I-III), atypical (HG IV-X), as well as HG XI and XII strains. IFNγ in the supernatant was measured by ELISA at 48h. Average of 2-12 experiments + SD for each strain is shown, each dot represents a single experiment. Statistical analysis was performed using one-way ANOVA with Bonferroni’s correction comparing the grouped average of all clade A strains against the grouped averages of clade B, C, D or F strains; * p ≤ 0.05 and ** p ≤ 0.01. **(B)** The clade A BOF strain, which encodes a lowly expressed single *ROP5B* gene (‘low’) and BOF complemented with an LC37 cosmid that expresses the entire clade A *ROP5* locus (‘A’) were assayed as described in Fig 1A. Average IFNγ detected in the supernatant at 48h of 3 experiments + SD is shown. Statistical analysis was performed by an unpaired two-tailed t-test; ** p ≤ 0.01.

### Activation and IFNγ differentiation of CD8 T cells are only partially inhibited by *T. gondii* ROP5

Following antigen-driven TCR stimulation (or ‘signal 1’), early activated T cells receive secondary cues from the environment including co-stimulation (‘signal 2’) and cytokines (‘signal 3’) to commit to the production of cytokines like IFNγ. Whether clade A strains, through ROP5 or other effectors, intersect one or several these activation steps to lower T57 IFNγ responses is unclear. To explore this issue further, we generated a ‘T-GREAT’ IFNγ reporter mouse line by crossing T57 with GREAT mice (85). GREAT mice report IFNγ transcription with an internal ribosomal entry site (IRES)-eYFP reporter cassette inserted between the stop codon and endogenous 3’UTR with the poly-A tail of the *Ifng* gene. Without abrogating translation of IFNγ, GREAT mice allow faithful detection of *Ifng* transcription via YFP fluorescence and flow cytometry (85). In addition, surface expression of the activation marker CD69 is a proxy for early TCR signaling events, and is one of the first markers expressed by naïve T lymphocytes after activation (86). In this way, the relative amount of TGD057 that has escaped the PV and ultimately presented by MHC 1 molecules can be inferred by T cell upregulation of CD69, and this can be measured independently of IFNγ transcription in T-GREAT cells. Naïve T-GREAT cells were co-cultured with BMDMs infected with RH (clade A), RH *Δrop5*, and ME49 (clade D) strains (Fig 6A), and the frequency of activated CD8 T cells (CD62L-CD69+ CD8+ T cells) (Fig 6B) and YFP levels (*Ifng*:YFP+ of CD62L-CD69+ CD8+ T cells) (Fig 6C) were measured by flow cytometry at 18 hours. Without parasite, few CD69+ or *Ifng*:YFP+ T-GREAT cells were detected in the co-culture, consistent with their naïve beginnings (Fig 6B-6C). In contrast and over a range of multiplicity of infections (MOIs), there was approximately three-fold more CD69+ CD8 T cells elicited by the ME49 compared to the RH strains (Fig 6B, 6D). Among T cells that have been activated (CD69+), a comparison of the *Ifng* transcript level revealed a six-fold increase in response to the ME49 compared to the RH strain (Fig 6C, 6E). Although removal of ROP5 enhances the activation and differentiation of T-GREAT cells to the RH *Δrop5* strain, *Ifng* transcript levels never equaled that elicited by ME49, especially at lower MOIs (Fig 6E). Therefore, the data show T57 cells do in-fact recognize the RH strain, but once activated this genetic background fails to elicit other signals necessary for full induction of the T57 IFNγ response. Moreover, the data implicate *T. gondii* genetic determinants other than ROP5 intersect CD8 T IFNγ responses at the activation and likely differentiation steps.

**Figure 6:**
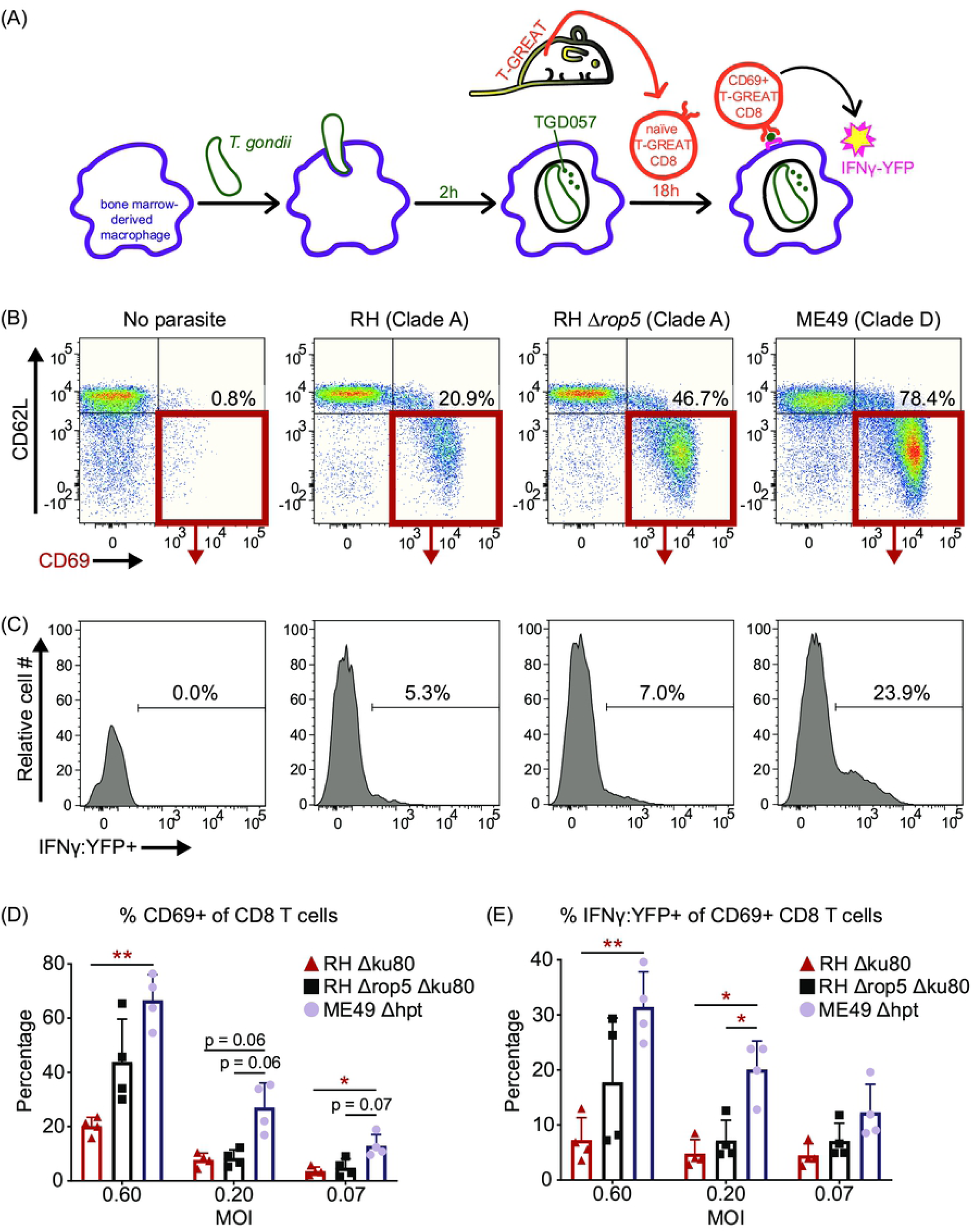
In the absence of ROP5, clade A strains still inhibit activation and IFNγ production by TGD057-specific CD8 T cells. **(A)** ‘T-GREAT’ reporter mice were generated by crossing T57 mice with an IFNγ (IRES)-eYFP reporter GREAT mouse line, which allows IFNγ transcript levels to be measured by flow cytometry as a function of YFP expression. T-GREAT cells were analyzed for activation (CD69+) and IFNγ-differentiation (*Ifng*:YFP+) in response to parasite-infected BMDMs at 18h. **(B-C)** The frequency of activated CD8 T cells (CD69+ CD62L-CD8+ T cells), as well as **(C)** the frequency of YFP+ (*Ifng*:YFP+) of activated CD69+ CD62L-CD8 T cells were compared between clade A RH, RH *Δrop5*, and clade D ME49 strains. Representative flow plots with indicated gates and percentages are shown. **(D)** Percent CD69+ CD62L-of total CD8 T cells, and **(E)** percent *Ifng*:YFP+ of CD69+ CD62L-CD8 T cells are shown. Each dot represents the results of an individual experiment and plotted is the average + SD of 4 experiments. Statistical analyses were performed using two-way ANOVA with Bonferroni corrections; * p ≤ 0.05, ** p ≤ 0.01.

### IFNγ-production by TGD057-specific CD8 T cells does not solely depend on IL-12

IL-12 signaling is essential for IFNγ-mediated control of *T. gondii* (87–89), and is required for full induction of IFNγ-producing KLRG1+ effector CD8 T cells following *T. gondii* type II infections in vivo (48, 49). Moreover, the RH strain fails to induce robust IL-12 secretion in infected macrophages (90, 91), perhaps underpinning the low T57 IFNγ responses to clade A strains observed in this system. To understand what extent IL-12 influences IFNγ-production, the T57 assay was performed with IL-12p40 deficient *Il12b-/-* BMDMs. The T57 IFNγ response to ME49 (Fig 7A), or MAS infected *Il12b-/-* BMDMs (Fig 7B) was reduced but not entirely abrogated compared to that of infected wildtype BMDMs. A partial reduction was also reported for adoptively transferred IL-12Rβ2-/- CD8 T cells that specifically lack IL-12 signaling during primary infection (48). Next, the T57 assay was performed with parasite strains known to regulate host IL-12 production. Three *T. gondii* effector proteins—GRA15, GRA24 and ROP16—modulate IL-12 production in infected BMDMs (92–95). Polymorphisms in GRA15 and ROP16 largely account for parasite strain differences in alternative (M2) and classical activation (M1) of macrophages (94). Specifically, polymorphisms in GRA15 render type II strains able to activate the NF-ĸB pathway through direct association with TRAF2 and TRAF6 (96), and its expression is an absolute requirement for IL-12p70 (94) and largely responsible for IL-12p40 production by type II-infected BMDMs (92). Through activation of host p38 MAPK, GRA24 promotes IL-12p40 and chemokine secretion by *T. gondii-*infected BMDMs (95). Although no consistent difference between parental type II (Pru) and GRA15-deficient or GRA24-deficient strains was observed, T57 IFNγ production was decreased but not abolished in response to a double deletion Pru *Δgra15 Δgra24* strain (Fig 7C). With respect to ROP16, in all *T. gondii* strains except those of clade D (97), the ROP16 kinase activates host STAT3, STAT5, STAT6 transcription factors (98–101), leading to the suppression of NF-ĸB signaling by an unknown mechanism (98, 101). When activating alleles of ROP16 are expressed as a transgene within the type II strain (Pru +*ROP16_A_*), it reduces IL-12 production in *T. gondii*-infected BMDMs and induces the expression of many M2 associated genes (94). The T57 IFNγ response to the Pru +*ROP16_A_* was reduced to half that of the parental strain (Fig 7D). Thus, although IL-12 and *T. gondii* GRA15, GRA24, and ROP16 have some impact in regulating TGD057-specific CD8 T cell IFNγ-production, the IL-12 axis does not fully account for IFNγ commitment in this system.

**Figure 7:**
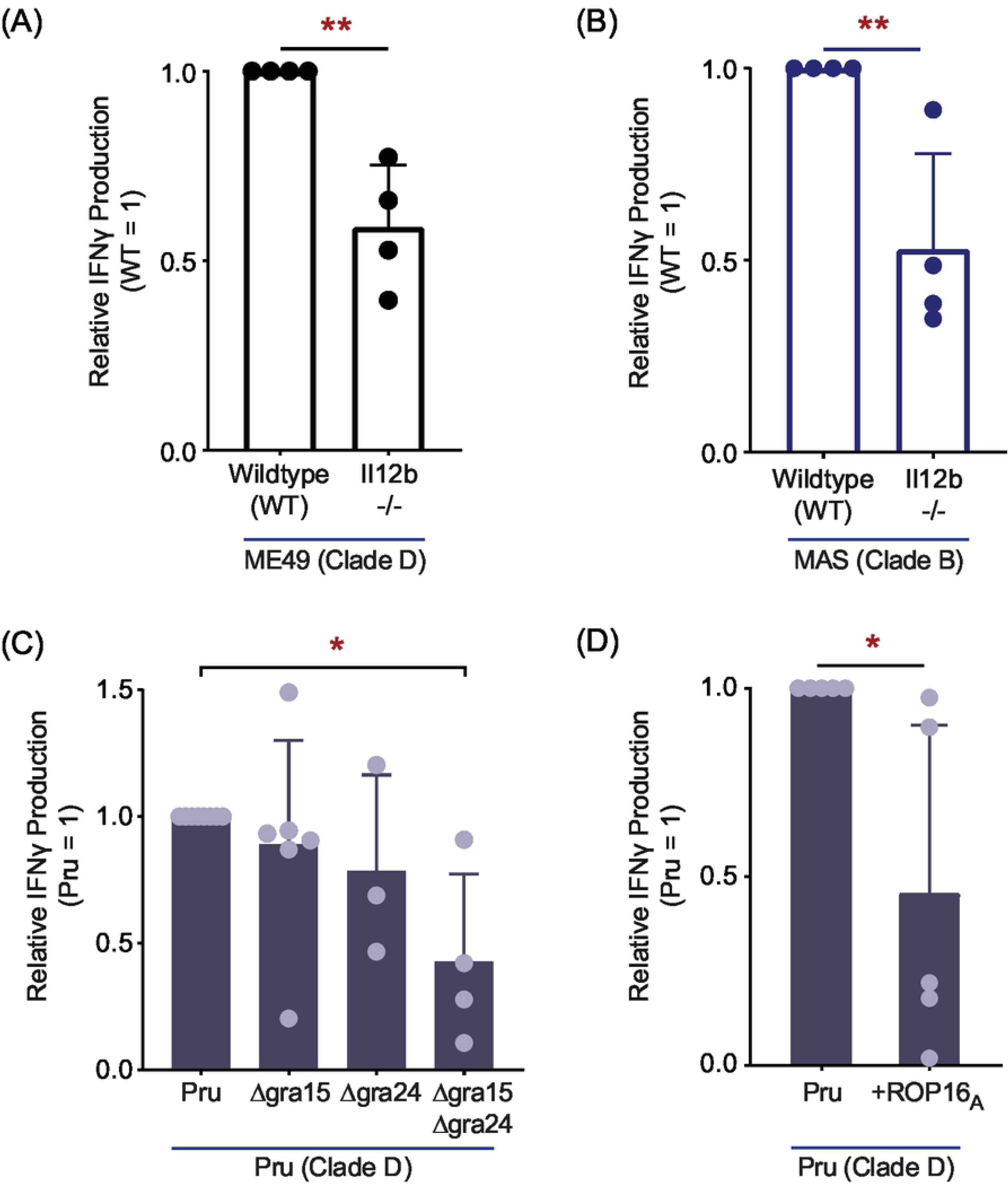
IL-12 signaling is partially required for TGD057-specifc CD8 T cell IFNγ responses. **(A)** *Il12b-/-* (IL-12p40) BMDMs were infected with clade D ME49, or **(B)** clade B MAS and TGD057-specific CD8 T cell IFNγ responses were measured as described in Fig 1A. Each dot represents the result of an individual experiment and the average of 4 experiments + SD is shown; statistical analysis was performed with an unpaired two-tailed t-test; ** p ≤ 0.01. **(C)** Various Pru (clade D) gene deletion strains, *Δgra15*, *Δgra24*, and *Δgra15*/*Δgra24*, or **(D)** a Pru strain transgenically expressing clade A ROP16 (Pru *+ROP16_A_*) were assayed for TGD057-specific CD8 T cell IFNγ responses. The IFNγ response, as analyzed by ELISA, is normalized to that induced by the wildtype Pru strain. Each dot represents the result of an individual experiment and the average of 3-6 experiments + SD is shown. Statistical analysis was performed using one-way ANOVA with Bonferroni’s correction for (C), and an unpaired two-tailed t-test for (D); * p ≤ 0.05.

### An inflammasome-independent NLRP3 pathway is required for maximal CD8 T cell IFNγ responses to *T. gondii*

Both NLRP3 and NLRP1 inflammasome activation occur following *T. gondii* infection (50,51,102), and IL-1 and IL-18 are known regulators of IFNγ production in a variety of cell types (103), including cells of the adaptive immune system (104). Therefore, BMDMs deficient at various steps in the inflammasome activation cascade were analyzed. In brief, most NLRP proteins undergo ASC-driven oligomerization, causing auto-activation of caspase-1, that in turn lead to the cleavage and maturation of IL-1β and IL-18 (105). Inflammasome activated caspase-1 and −11 also activate Gasdermin D, a key pore forming protein responsible for pyroptosis and extracellular release of IL-1/18 in several biological contexts (106–111). The T57 IFNγ-response was largely reduced to parasite-infected *Nlrp1-/-* BMDMs, and completely absent to infected *Nlrp3-/-* BMDMs (Fig 8A-8B). In contrast, the IFNγ response was only partially decreased to *Asc-/-*, *Casp1/11-/-*, and *Gsdmd-/-* BMDMs infected with ME49 (Fig 8A), and no consistent difference was observed between knockout and wildtype BMDMs infected with MAS (Fig 8B). These results indicate CD8 T cell IFNγ differentiation, though entirely dependent on NLRs, only partially involves inflammasome matured IL-1 and/or IL-18. Consistent with this supposition, the CD8 T cell IFNγ response to parasite-infected macrophages is reduced but not abrogated in the absence of IL-18 and IL-1R-signaling, as assessed with neutralizing antibodies that block IL-1β and IL-1α engagement with its receptor, IL-1R, and *Il18*-/- BMDMs (Fig 8C). Finally, to test whether NLRP3 impacts T cell activation or differentiation, T-GREAT CD8 T cell activation profiles to parasite-infected *Nlrp3*-/- BMDMs were explored. Whereas the percentage of early activated CD69+ T-GREAT cells in response to ME49 and MAS infections was slightly decreased to *Nlrp3-/-* BMDMs compared to wildtype BMDMs (Fig 9A), IFNγ transcript levels in the activated CD69+ population dropped by 50-70% (Fig 9B), suggesting NLPR3 induces a macrophage-derived signal required for IFNγ transcription in activated CD8 T cells. To summarize, although inflammasome matured cytokines may play some role in promoting T57 IFNγ production, there is no substitute for NLRP3, and to a similar extent NLRP1 in this system. Importantly, our data identify a novel NLR-dependent but NLR-ASC inflammasome complex-independent pathway that regulates CD8 T cell IFNγ responses to an intracellular pathogen.

**Figure 8:**
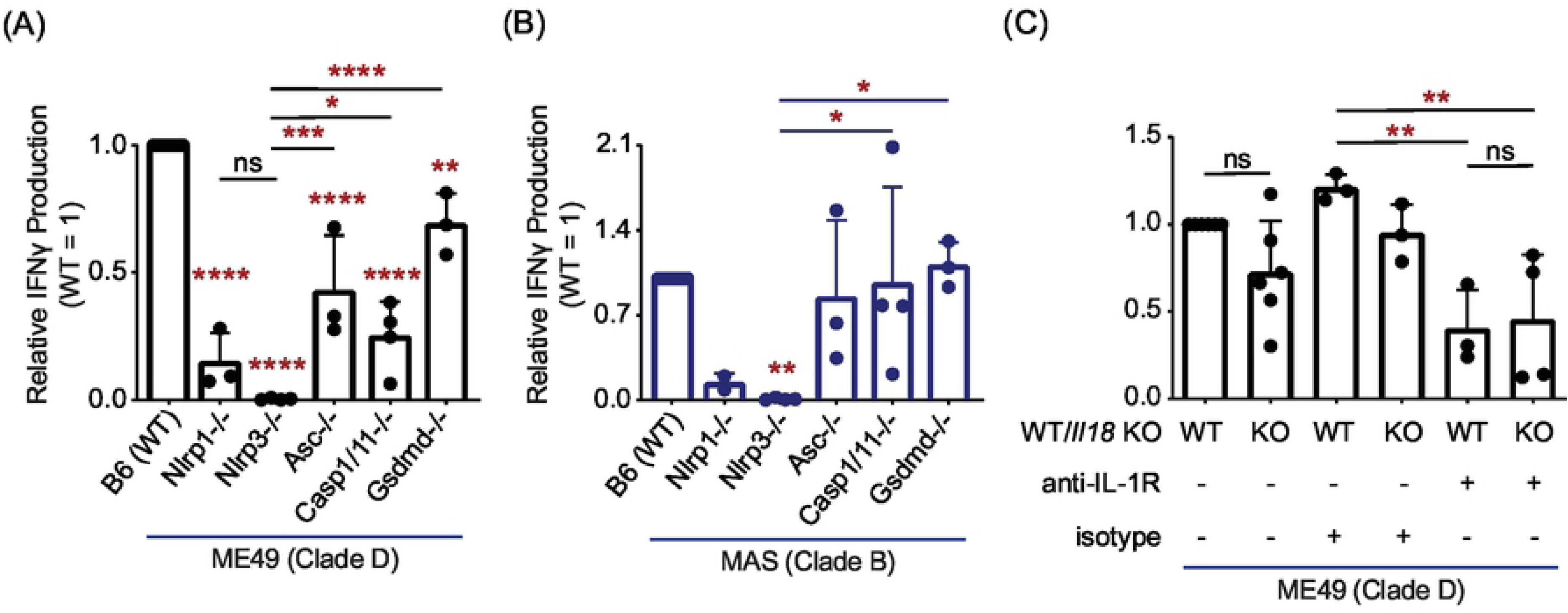
An inflammasome-independent NLR pathway promotes CD8 T cell IFNγ responses to *T. gondii.* **(A)** BMDMs with indicated gene deletion were infected with the clade D ME49, or **(B)** clade B MAS strain. The T57 CD8 T cell IFNγ response to TGD057 was analyzed by ELISA and normalized to that of WT BMDMs. Each dot represents the result of an individual experiment, and the average of 2-4 experiments + SD is shown. Statistical analysis was performed using one-way ANOVA with Bonferroni’s corrections compared to WT or *Nlrp3*-/-, the latter comparisons are indicated with a line; * p ≤ 0.05, ** p ≤ 0.01, *** p ≤ 0.001, **** p ≤ 0.0001, ns non-significant. **(C)** WT and *Il18*-/- BMDMs were infected with the clade D ME49 and assayed for CD8 T cell T57 IFNγ production. Additionally, co-cultures were treated with either anti-IL-1R neutralization or isotype control antibodies. The IFNγ level was measured by ELISA and normalized to that of untreated WT BMDMs. Each dot represents the results from an individual experiment and the average of 3-6 experiments + SD is shown. Statistical analysis was performed with one-way ANOVA and Bonferroni’s corrections compared to untreated WT BMDMs; ** p ≤ 0.01, ns non-significant.

**Figure 9:**
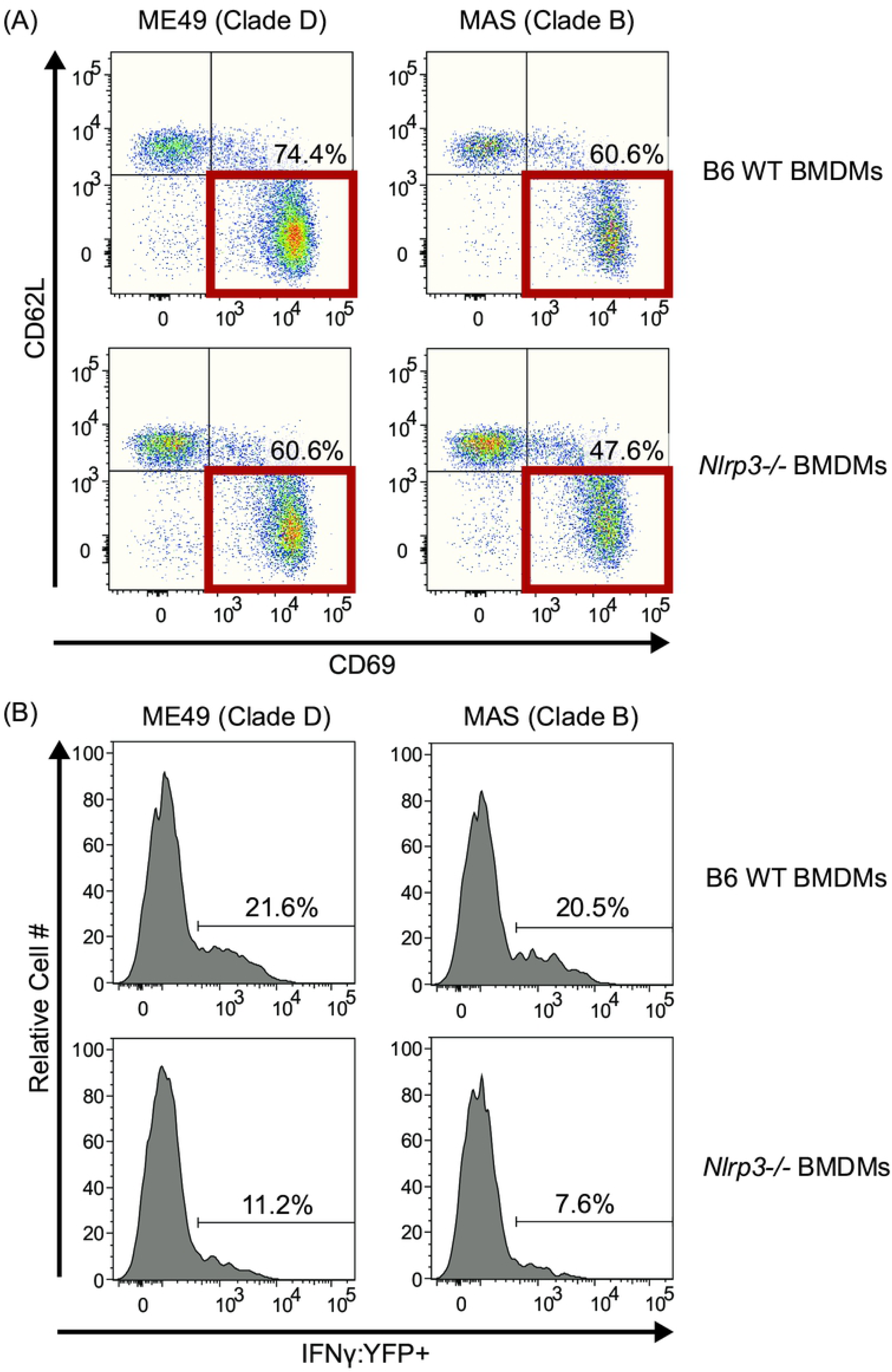
IFNγ transcription in activated CD8 T cells is promoted by NLRP3 in *T. gondii*-infected cells. WT and *Nlrp3-/-* BMDMs were infected with clade D ME49 or clade B MAS strains. T-GREAT CD8 T cell responses to parasite-infected BMDMs were analyzed as described in Fig 6A. **(A)** The frequency of activated CD8 T cells (CD69+ CD62L-CD8+ T cells), and **(B)** the frequency of YFP+ (*Ifng*:YFP+) of activated CD69+ CD62L-CD8 T cells were determined. Representative flow plots with indicated gates and frequencies are shown from 2-3 experiments.

## DISCUSSION

In this work, we set out to explore the role of parasite virulence factors, and host pathways that regulate CD8 T cell IFNγ responses to an endogenous antigen. Given the close association between T cell activation and parasite-infected cells *in vivo* (33), and the immune pressure incurred by CD8 T cells, we reasoned there may be unidentified mechanisms governing host-pathogen interactions between *T. gondii* and this cell type. To this end, we used T57 transnuclear mice to develop a system to study this interaction, which has several advantages including the conserved nature of the TGD057 epitope and the ability to readily analyze responses of antigen-specific clonal CD8 T cell population with a normally expressed TCR. Without need for further antigen cloning, certain matters surrounding CD8 T cell IFNγ differentiation and MHC 1 antigen presentation of TGD057 were revealed, including a fundamental role for *T. gondii* ROP5 and host NLRP3 in regulating this response. Our observations are both similar and divergent from results obtained in other systems, pointing to the contextual nature of immune responses to live pathogens, but also to new host mechanisms that possibly promote CD8 T cell immunity to *T. gondii*.

First, many similarities exist between the CD8 T cell response to TGD057 and OVA. For example, CD8 T cell activation to both antigens utilize the host’s IRG system for optimal responses (30,44,45). Such results indicate a need for the host to actively acquire antigens sequestered inside a PV (39). We lend credence to this hypothesis, in that the parasite’s export machinery appears dispensable for inducing TGD057-specific CD8 T cell responses (Fig 2). One curious observation in this regard, is that the TGD057 peptide epitope, SVLAF<u>RRL</u>, encodes the lone ASP5 recognition TEXEL motif (underlined) (Fig S1). In fact, ASP5 cleavage would preferentially produce parasite peptides with a terminal leucine, which is the preferred anchor residue of the P8/9 peptide binding pocket of MHC 1 K^b^ (112) and many other murine MHC and human HLA alleles. We initially thought ASP5 might prepare *T. gondii* antigens for binding host MHC 1 molecules, such that in the absence of ASP5, a blunted CD8 T cell response would ensue. Rather, the opposite occurred (Fig 2), ruling against parasite-assisted antigen processing of TGD057. In addition to its role in protein export, ASP5 is required for targeting dense granules to the PVM (59,60,113). GRA6 association with the PVM significantly enhances its entry into the host’s MHC 1 antigen presentation pathway (41). Whether TGD057 sub-localization inside the PV impacts entry into host MHC 1 antigen presentation pathways is currently unknown. The enhanced response to the type II *Δasp5* strain (Fig 2) may also reflect decreased parasite fitness observed for this strain (59), or the role of an unidentified ASP5-targeted and PVM-associated GRA that inhibits CD8 T cell activation. Such possibilities await experimental validation.

Second, since *T. gondii* defends itself from immune attack by its virulence factor ROP5, this strategy affords a second benefit, the hiding of its vacuolar antigens from the host’s MHC 1 antigen processing machinery. For TGD057, this battle is uniquely defined by ROP5A that has no known interacting partner, but suppresses the CD8 T cell response to TGD057. We predict this occurs by a mechanism independent from any known ROP5-IRG interaction. For example, ROP5B and ROP5C from clade A strains are able to prevent accumulation of effector IRGs on the parasite’s PVM, including Irgb10, Irga6 and Irgb6, but this is not a function of clade A ROP5A (3, 4), nor is this a known function for any ROP5 isoforms from clade D. Although ROP5C was not tested here, recently an OVA-expressing RH *Δrop5* +*ROP5C* strain was generated and CD8 T cell activation was partially inhibited by this strain (45). Whatever mechanism underlies the ability of ROP5 to inhibit CD8 T cell activation, we hypothesize that it is controlled by ROP5A and ROP5B isoforms, and less so by ROP5C. Amino-acids in the 346-370 region in the IRG binding interface of ROP5 (3) can be found that distinguish clade A ROP5C from ROP5A and ROP5B of clades A and D (not shown). Whether these amino acids define a novel interaction with a less-studied effector IRGs or another host protein is currently unknown.

It is likely there is no single host-parasite interaction that determines antigen presentation for all *T. gondii* vacuolar antigens. For example, one notable difference between CD8 T cell responses to the vacuolar TGD057 and OVA antigens, is the role of the *T. gondii* ROP18 kinase. The CD8 T cell response to OVA appears largely inhibited by ROP18 (45), but at best, ROP18 plays a marginal role in this system (Fig 4, S3A). These observations imply effector IRG modulation or the ATF6β ER-stress response, which is known to regulate CD8 T cell IFNγ responses to *T. gondii* and is directly antagonized by the kinase activity of ROP18 (114), might be more important for CD8 T cell detection of OVA than TGD057. Furthermore, p62 is required for OVA-specific OT1 CD8 T cell activation by a mechanism that includes binding to ubiquitin-tagged PVs in IFNγ-stimulated cells (44). P62 recruits GBPs to the vacuole of *T. gondii* (115), suggesting GBPs may also assist in the MHC 1 antigen presentation, for which we found some evidence (Fig 3). Although the role of p62 using mouse knockout cells was not directly tested, a comparison of p62-PV localization patterns between stimulatory and non-stimulatory parasites strains (Fig S4) argues against a dominant role for this pathway in our system. Other differences include lessons learned from the GRA6 antigen. When the C-terminal GRA6 epitope is facing the host cytosol it is highly stimulatory to CD8 T cells (42). The protruding nature of the GRA6 epitope into the host cytosol may bypass need for host recruitment of IFNγ-induced IRG/GBP machinery, thus facilitating its immuno-dominance. However, TGD057 is not an integral membrane protein nor is it associated with the membranous fractions of the PV (41). It is therefore unclear how the initial antigen is first detected to start T57 IFNγ responses, which paradoxically require IFNγ signaling to begin with (Fig 3). A clue may come from the OVA system. Host derived ER-GICs fuse with the PVM in a Sec22b SNARE-dependent process to initiate MHC 1 presentation of *T. gondii* expressed OVA (43). Whether this pathway seeds the initial antigen-specific response to TGD057 is unknown. Yet even in response to clade A strains, which are poor inducers of TGD057-specific CD8 T cell IFNγ and IL-2 responses (Fig 5, S3B), the early activation marker CD69 was readily detected on T57 CD8 T cells (Fig 6). The immune system is therefore robust in its ability to perceive *T. gondii* antigens, which employs multiple non-redundant pathways to acquire antigens from vacuolated pathogens (116).

Third, our studies demonstrate an absolute requirement for the pathogen sensor NLRP3, and to a similar extent NLRP1, for promoting naïve TGD057-specific CD8 T cell IFNγ responses to parasite-infected cells. Moreover, it appears NLR-mediated regulation can occur in the absence of other components of the inflammasome cascade. This is inferred because T57 IFNγ production was still detected in response to parasite-infected ASC, caspase-1/11, and gasdermin D deficient cells, or when IL-1/18 cytokine signaling was inhibited. In contrast, when NLRP3 is removed, there was no IFNγ response (Fig 8), even though the CD8 T cells were activated (Fig 9). NLRs have several inflammasome independent functions, including the ability to form a bridge between ER and mitochondria to initiate inflammasome signaling (117). NLRs also bind to and directly activate transcription factors, such as IRF4 (118) and CIITA, the latter induces MHC expression (119). Our results diverge from a recent report exploring the role of inflammasome components in promoting CD8 T cell IFNγ responses during primary *T. gondii* infection. NLRP3, ASC and caspase 1/11 deficiency had no bearing on the frequency of peritoneal or splenic IFNγ+ CD8 T cells during primary *T. gondii* infection (52). Certainly, CD8 T cells receive environmental cues *in vivo* that compensate for the lack of the inflammasome pathway, but are missing in our system. Given the diminished IFNγ-transcription response to NLRP3-deficient cells (Fig 9), it is possible that the missing signal is co-stimulation. Co-stimulation determines post-transcriptional regulation of IFNγ in tumor-infiltrating T cells (120), and in its absence, enforces post-transcriptional silencing of IFNγ in anergic self-reactive T cells (121). Current studies are underway to understand the mechanism by which NLRP3 deficiency impacts transcriptional and translation regulation of IFNγ in activated CD8 T cells.

Finally, we present evidence that *T. gondii* strains differ in their ability to modulate CD8 T cell IFNγ responses, which in theory might aid the parasite’s survival in a broad host range. For example, T57 IFNγ cell responses were low to *ROP5*-expressing clade A strains, which we initially hypothesized might be due to *ROP5* polymorphisms unique to clade A strains. However, both clade D and A *ROP5* alleles were equally able to repress the CD8 T cell IFNγ response (Fig 4), indicating another polymorphic regulator is in effect. Our current work searching for this polymorphic modulator of host CD8 T cell IFNγ response, has revealed no genotype-phenotype correlation at the *ROP5* locus (not shown). Instead, the polymorphic regulator could intersect CD8 T cell differentiation, for example through modulation of the host’s NLRs, or antigen release, possibly by assisting the function of ROP5A. Recently, Rommereim *et al.* has shown OVA-specific CD8 T cell activation is regulated by multiple GRA(s), including those that modulate *T. gondii’s* intravacuolar network (IVN) inside the PV (45). Whether any such GRAs are responsible for the strain differences in T57 activation require further investigations.

In summary, since any warm-blooded animal can serve as an intermediate host for *T. gondii*, the parasite may have difficulty achieving stable chronic infections in every animal to promote its transmission. This is evidenced by the observation that *T. gondii* strains differ dramatically in virulence in laboratory mice (122) and correlate with the severity of human toxoplasmosis (123–126). This led to the hypothesis that parasite strains have adapted to certain intermediate host niches (127), defined by host genetics, including that of the murine IRG locus (128). This adaptation may also necessitate the manipulation of host adaptive immune responses. Here, we present evidence that *T. gondii* may direct CD8 T cell IFNγ response for its advantage. Perhaps NLRP3 and the MHC 1 antigen presentation pathway serve as two distinct sites for immune pressure, leading to the evolution of novel parasite virulence factors. Nevertheless, a closer and detailed understanding of the interactions between *T. gondii* and host CD8 T cells will eventually help us to find potential therapeutic targets for toxoplasmosis, as well as to understand why *T. gondii* has spread so extensively.

## MATERIALS AND METHODS

### Ethics statement

All animal protocols were approved by UC Merced’s Committee on Institutional Animal Care and Use Committee (IACUC) (AUP17-0013). All mouse work was performed in accordance with the recommendations in the *Guide to the Care and Use of Laboratory Animals* of the National Institutes of Health and the Animal Welfare Act (assurance number A4561-1). Inhalation of CO_2_ to effect of 1.8 liters per minute was used for euthanasia of mice.

### Parasites

Tachyzoites of *Toxoplasma gondii* strains were passaged in ‘Toxo medium’ [4.5 g/liter D-glucose in DMEM with GlutaMAX (Gibco, cat#10566024), 1% heat-inactivated fetal bovine serum (FBS) (Omega Scientific, cat#FB-11, lot#441164), 1% penicillin-streptomycin (Gibco, cat#15140122)], in confluent monolayers of human foreskin fibroblasts (HFFs). HFFs were cultured in ‘HFF medium’ [4.5 g/liter D-glucose in DMEM with GlutaMAX (Gibco), 20% heat-inactivated FBS (Omega Scientific), 1% penicillin-streptomycin (Gibco), 0.2% Gentamicin (Gibco, cat#15710072), 1X L-Glutamine (Gibco, cat#21051024)]. Strains assayed include GT1 (type I, clade A), BOF (HG VI, clade A), FOU (HG VI, clade A), CAST (HG VII, clade A), MAS (HG IV, clade B), TgCatBr5 (HG VIII, clade B), CEP *hxgprt-* (type III, clade C), P89 (HG IX, clade C), ME49 *Δhxgprt::Luc* (129) (type II, clade D), Pru *Δhxgprt* (type II, clade D), Cougar (HG XI, clade D), B73 (HG XII, clade D), GUY-KOE (HG V, clade F), GUY-MAT (HG V, clade F), RUB (HG V, clade F), GUY-DOS (HG X, clade F), and VAND (HG X, clade F). Other strains used include BOF+LC37 (75), RH *Δhxgprt* (130), RH *Δhxgprt Δku80* (131), RH *Δhxgprt Δku80 Δrop5::HXGPRT* (RH *Δrop5*) (9), RH *Δhxgprt Δku80 Δtgd057::HXGPRT* (RH *Δtgd057*) (26), RH *Δhxgprt Δku80 TGD057-HA::HXGPRT* (RH_tgd057_-HA) (generated here), Pru *Δhxgprt Δku80,* Pru *Δhxgprt Δku80 Δtgd057::HXGPRT* (Pru *Δtgd057*) (generated here), Pru A7 Δ*hxgprt::gra2-GFP::tub1-FLUC* (Pru A7) (132), Pru A7 *Δhxgprt Δgra15::HXGPRT* (Pru *Δgra15*) (92), Pru A7 *Δhxgprt Δgra24* (1E9) (Pru *Δgra24)* (generated here), Pru *Δhxgprt Δgra15 Δgra24* (1F8) (Pru *Δgra15 Δgra24*) (generated here), Pru A7 *Δhxgprt +ROP16_A_::HXGPRT* (Pru +*ROP16_A_*) (94), RH *Δhxgprt Δmyr1::HXGPRT* (RH *Δmyr1*) (113), RH *Δku80 Δasp5-ty::DHFR* (RH *Δasp5*) (65). ME49 *Δasp5::DHFR* (ME49 *Δasp5*) was a generous gift from Dominique Soldati-Favre (University of Geneva) (59). ME49 *Δhxgprt::Luc Δmyr1::HXGPRT* (ME49 *Δmyr1*), ME49 *Δhxgprt::Luc Δmyr1 +MYR1-HA::HXGPRT* (ME49 *Δmyr1::MYR1*) (61), and RH *Δhxgprt Δrop17::HXGPRT* (PCRE) (RH *Δrop17*) (133) were generous gifts from John Boothroyd (Stanford University). The ROP5 complementation strains used in this study are as follows: RH *Δhxgprt Δku80 Δrop5* +*ROP5A_II-ME49_His6-3xFlag* (C1A6) (RH *Δrop5 +ROP5A_D_*) (generated here), RH *Δhxgprt Δku80 Δrop5* +*ROP5B_III-CTG_-His6-3xFlag* (H2) (RH *Δrop5 +ROP5B*) (generated here), RH *Δhxgprt Δku80 Δrop5* +*ROP5A_III-CTG_His6-3xFlag* (C1B1) (RH *Δrop5 + ROP5A)*, RH *Δhxgprt Δku80 Δrop5 Δuprt::ROP5A_III_-HA +ROP5B_III-CTG_-His6-3xFlag* (AC11) (RH *Δrop5 +ROP5A+ROP5B)*, RH *Δhxgprt Δku80 Δrop5 Δuprt::ROP5A_III_-HA +ROP5A_III-CTG_-His6-3xFlag* (AA12) (RH *Δrop5 +ROP5A+ROP5A*) (9).

### Generation of gene knock-in and knockout parasite strains

The Pru *Δhxgprt Δku80 Δtgd057::HXGPRT* strain was generated using the same primers and strategy as previously described (26). The RH *Δhxgprt Δku80 TGD057-HA::HXGPRT* endotagged strain was generated as previously described (134). In brief, for endogenous tagging (131) of TGD057 with an HA tag, the gene (*TGGT1_215980*) was amplified with a forward primer internal to the ATG start site of *TGD057* [“AC_tgd057endoF3” 5’-CACCAACTACGTCGGAGCGCCTGTACG-3’], containing a 5′-CACC-3′ sequence required for directional TOPO cloning in pENTR/D-TOPO (Invitrogen, USA), and a reverse primer [“Tgd057 R HA stop” 5’-TTA<u>CGCGTAGTCCGGGACGTCGTACGGGTA</u>CTCGACCTCAATGTTGTATTC-3’], containing the hemagglutinin (HA) tag sequence (underlined) followed by a stop codon. The resulting TGD057 HA-tagged DNA fragment was then cloned into the pTKO-att parasite expression vector (92) by Gateway Recombination Cloning Technology (Invitrogen). The resulting vector was linearized and transfected into RH *Δhxgprt Δku80* parasites by electroporation in a 2 mm cuvette (Bio-Rad Laboratories, USA) with 2 mM ATP (MP Biomedicals) and 5 mM glutathione (EMD) in a Gene Pulser Xcell (Bio-Rad Laboratories), with the following settings: 25 μFD, 1.25 kV, ∞ Ω. Stable integrants were selected in media with 50 μg/ml of mycophenolic acid (Axxora) and 50 μg/ml of xanthine (Alfa Aesar) and cloned by limiting dilution. The correct tagging was confirmed by PCR, using a primer upstream of the plasmid integration site and a primer specific for the HA tag (5’-CGCGTAGTCCGGGACGTCGTACGGGTA-3’), and by an immunofluorescence assay (IFA) using an HA-specific antibody (Sigma, clone 3F10).

For the generation of RH *Δrop5 +ROP5A_D_* and RH *Δrop5 +ROP5B* complementation strains, RH *Δhxgprt Δku80 Δrop5::HXGPRT* tachyzoites were transfected by electroporation with linearized pTKO-“ROP5KO” -*ROP5A_II-ME49_-His6-3xFlag* plasmid (9), or a similarly generated pTKO -*ROP5B_III-CTG_-His6-3xFlag* plasmid as described in (9). In brief, for both strains the complementation allele, *ROP5-His6-3xFlag*, is flanked by homology arms to the *Δrop5::HXGPRT* locus, whereby the transfected population is selected for removal of *HXGPRT* and replacement with the complementation allele in 6-thioxanthine selection medium [177 µg/mL of 6-thioxanthine (TRC, cat# T385800) in 4.5 g/liter D-glucose in GlutaMAX DMEM (Gibco), with 1% dialyzed FBS (Omega Scientific, cat#FB-03, lot#463304)]. Post-selection and limiting dilution cloning, *ROP5-His6-3xFlag* complementation strains were assessed by IFA with a mouse anti-Flag primary antibody (Sigma, clone M2) at 1:500 dilution and Alexa Flour 594 goat anti-mouse IgG secondary antibody (Life Technologies) at 1:3000 dilution. Both clones expressed the ROP5-HF in the expected rhoptry organelles by IFA (not shown).

For generating the Pru A7 *Δhxgprt Δgra24* strain, Pru A7 Δ*hxgprt* parasites were transfected with a NotI-linearized plasmid expressing a loxP-flanked pyrimethamine selectable cassette (*loxP-DHFR-mCherry-loxP*) (Addgene plasmid #70147, was a gift from David Sibley, Washington University in St. Louis) and a CRISPR-CAS9 construct targeting *GRA24* (*TGME49_230180*). Transfectants were selected and cloned in medium containing pyrimethamine, and screened for the disruption of *GRA24*. For generating the Pru A7 *Δhxgprt Δgra15 Δgra24* strain, first the Pru A7 *Δhxgprt Δgra15::HXGPRT* strain (92) was gene-edited with CRISPR-CAS9 targeting *HXGPRT* and selected against its expression with 6-thioxanthine. Then, a Pru A7 *Δhxgprt Δgra15::hxgprt-* clone was used to make a Pru A7 *Δhxgprt Δgra15 Δgra24* double knockout strain using the same method as described above. Finally, the *loxP-DHFR-mCherry-loxP* cassette was removed from both *Δgra24* and *Δgra15*/*Δgra24* strains by transfection with a Cre recombinase parasite-expression plasmid (135), and immediately cloned by limiting dilution. The details of which can be found in a later manuscript by Mukhopadhyay et al.

### Immunofluorescence Assay

For TGD057 visualization, HFFs were seeded on coverslips with HFF medium in 24-well tissue culture-treated plates. The confluent monolayer HFFs were infected with *T. gondii* and incubated at 37°C, 5% CO_2_ overnight. For Irgb6- and p62-PV localization, BMDMs or mouse embryonic fibroblasts (MEF) were plated on coverslips with ‘BMDM medium’ [4.5 g/liter D-glucose in DMEM with GlutaMAX (Gibco), 20% heat-inactivated FBS (Omega Scientific), 1% penicillin-streptomycin (Gibco), 1X non-essential amino acids (Gibco, cat#11140076), 1mM sodium pyruvate (Gibco, cat#11360070)] supplemented with 20% L929 conditioned medium or ‘MEF medium’ [4.5 g/liter D-glucose in DMEM with GlutaMAX (Gibco), 20% heat-inactivated FBS (Omega Scientific), 1% penicillin-streptomycin (Gibco), 0.2% Gentamicin (Gibco), 20 mM HEPES (Gibco, cat#15630080)], respectively, in 24-well tissue culture-treated plates. The BMDMs and MEFs were treated with 20 ng/ml of IFNγ overnight. The cells were then infected with *T. gondii* and incubated at 37°C, 5% CO_2_ for 3-4 hours. The samples were fixed with 3% formaldehyde in phosphate buffered saline (PBS) for 20 minutes and blocked with blocking buffer (3% BSA, 5% normal goat serum or fetal bovine serum depending the species of antibody used, 0.2% Triton X-100, 0.1% sodium azide in PBS). To visualize TGD057-HA, RH or RH_tgd057_-HA were stained with rat anti-HA primary antibody (Sigma, clone 3F10) at 1:500 dilution, followed by Alexa Fluor 594 goat anti-rat IgG (Life Technologies) secondary antibody (1:3000 dilution). To visualize p62, infected cells were stained with mouse anti-p62 (anti-SQSTM1) primary monoclonal antibody (Abnova, clone 2C11) at 1:50 or 1:100 dilutions, followed by Alexa Fluor 594 goat anti-mouse IgG (Life Technologies) secondary antibody at 1:3000 dilution. To visualize Irgb6, infected BMDMs were stained with TGTP goat polyclonal primary antibody (Santa Cruz Biotechnology, sc-11079) at 1:100 dilution, followed by Alexa Fluor 594 donkey anti-goat IgG (Life Technologies) at 1:3000 dilution. *T. gondii* PVM was stained with polyclonal rabbit anti-GRA7 primary antibody (gift from John Boothroyd, Stanford University) and Alexa Fluor 488 anti-rabbit IgG (Life Technologies) at 1:3000 dilution. Host nuclei were visualized with DAPI (Thermo Fisher, cat#62248) at 1:10,000 dilution or Hoechst (Life Technologies, cat# H3075) at 1:3000 dilution.

### Mice and generation of bone marrow-derived macrophages

Six-week-old female *Stat1-/-* (colony 012606), *Ifngr-/-* (colony 003288), *Nlrp1-/-* (colony 021301), *Nlrp3-/-* (colony 021302), *Casp1/11-/-* (colony 016621), *Il18-/-* (colony 004130), and *Il12b-/-* (colony 002693) and wildtype C57BL/6J (B6) (colony 000664) mice were purchased from Jackson Laboratories, and all of the C57BL/6 background. C57BL/6 *Asc-/-* mice were generous gifts from Vishva Dixit (Genentech). Hind bones from C57BL/6 *Gsdmd-/-* mice (136) were generous gifts from Igor Brodsky (University of Pennsylvania). *Irgm1-/-* and *Irgm1/m3-/-* hind bones were provided from Gregory Taylor (Duke University). *GBP^chr3^-/-* hind bone marrow cells were provided by Masahiro Yamamoto (Osaka University). Bone marrow cells were obtained and cultured in BMDM medium supplemented with 20% L929 conditioned medium. After 6-7 days of differentiation, BMDMs were harvested and were 98% pure CD11b+ CD11c-macrophages by FACS (not shown). *Asc-/-*, *Gsdmd-/-*, and *Nlrp3*-/- BMDMs were stained with anti-mouse MHC 1 K^b^-PE labeled antibodies (BioLegend, clone AF6-88.5) and they were positive by FACS analysis.

Transnuclear T57 mice (53) were bred in-house under specific pathogen free (SPF) conditions. ‘T-GREAT’ mice were generated by back- and inter-crossing between T57 and IFNγ-stop-IRES:eYFP-endogenous poly-A tail reporter mice (GREAT mice) (85), such that breeders obtained from F3 intercrossed mice were homozygous at three alleles: T57 TCRα (*TRAV6-4 TRAJ13* rearrangement), T57 TCRβ (*TRBV13-1 TRBJ2-7* rearrangement), and the GREAT reporter. T-GREAT mice were then maintained in our SPF facility with no overt fitness defects observed. Genotyping primers were as followed: GREAT allele (FW 5’-CCATGGTGAGCAAGGGCGAGG-3’; RV 5’-TTACTTGTACAGCTCGTCCAT-3’); wildtype *Ifng* allele (FW 5’-CAGGAAGCGGAAAAGGAGTCG-3’; RV 5’-GTCACTGCAGCTCTGAATGTT-3’); T57 TCRα (*TRAV6-4 TRAJ13* rearrangement: “146-alpha” FW 5’- GATAAGGGATGCTTCAATCTGATGG-3’; “108-alpha” RV 5’- CTTCCTTAGCTCACTTACCAGGGCTTAC-3’); endogenous non-rearranged *TRAV6-4* and *TRAJ13* loci (“191-alpha” FW 5’-GAGGCTTTACGTTAGTGATCTAAAC-3’; “108-alpha” RV); T57 TCRβ (*TRBV13-1 TRBJ2-7* rearrangement: “91-beta” FW 5’-CTTGGTCGCGAGATGGGCTCCAG-3’; “103-beta” RV 5’-GTGGAAGCGAGAGATGTGAATCTTAC-3’); endogenous non-rearranged *TCRBV13-1* and *TRBJ2-7* loci (“142-beta” FW 5’-GCACTCGGCTCCTCGTGTTAGGTG-3’; “103-beta” RV).

### T cell activation assay

2×10^5^ BMDMs cells were plated per well in a 96-well tissue culture-treated plate, in BMDM medium supplemented with 10% L929 conditioned medium. The following day, these BMDMs were infected with *T. gondii* tachyzoites in T cell medium (RPMI 1640 with GlutaMAX (Gibco, cat#61870127), 20% heat-inactivated FBS (Omega Scientific, cat#FB-11, lot#441164), 1% penicillin-streptomycin (Gibco, cat#15140122), 1 mM sodium pyruvate (Gibco, cat#11360070), 10 mM HEPES (Gibco), 1.75 µl of β-mercaptoethanol (Gibco, cat#21985023) per 500 mL RPMI 1640 with GlutaMAX. The infections were performed in triplicates, at MOI 0.6, 0.2, and 0.07. Then, lymph nodes and spleens were obtained from either T57 or T-GREAT transnuclear mice. The lymph node cells and splenocytes were combined and red blood cells were lysed with ammonium chloride-potassium (ACK) lysis buffer. 5×10^5^ cells were added into each well of the infected BMDMs (approximately 2 hours post-infection). For IL-1R neutralization, 50 µg/mL of anti-mouse IL-1R antibody (BioXCell, clone JAMA-147) or 50 µg/mL of isotype control (BioXCell, cat#BE0091) were added when BMDMs were infected.

### Correction for relative viability between parasites

Confluent monolayer HFFs, seeded in 24-well plates, were infected with 100 and 300 parasites. Plaques were counted 4-6 days after infection. Displayed results are from MOIs with similar viability, the equivalent of ∼MOI 0.2 was chosen for most assays.

### ELISA

The concentration of cytokines in the 24h and 48h supernatants from the T57 T cell activation assay was measured by ELISA according to the manufacturer’s instructions (IFNγ: Invitrogen eBioscience, cat#88731477, IL-2: Invitrogen eBioscience, cat#88702477, IL-17A: Invitrogen eBioscience, cat# 88737188). The supernatants were analyzed at various dilutions (1:2, 1:20, and 1:200) to obtain values within the linear range of the manufacture’s ELISA standards.

### Flow cytometry

At 18h after T57 T cell activation, samples were harvested for FACS analysis. With preparations all done on ice, cells were washed with FACS buffer [PBS pH 7.4 (Gibco, cat#10010049), 2% heat-inactivated FBS (Omega Scientific)] and blocked with blocking buffer [FACS buffer with 5% normal Syrian hamster serum (Jackson Immunoresearch, cat#007-000-120), 5% normal mouse serum (Jackson Immunoresearch, cat#015-000-120), and anti-mouse CD16/CD32 FcBlock (BD Biosciences, clone 2.4G2) at 1:100 dilution)]. Then, the samples were stained with fluorophore-conjugated monoclonal antibodies at 1:120 dilution – anti-mouse CD8α PE (eBioscience, clone 53-6.7), anti-mouse CD3ε APC-eFlour780 (eBioscience, clone 17A2), anti-mouse CD62L eFlour450 (eBioscience, clone MEL-14), and anti-mouse CD69 APC (BioLegend, clone H1.2F3). Samples were then stained with propidium iodide (PI) at 1:1000 dilution (Sigma, cat#P4170). Flow cytometry was performed on an LSRII (Becton Dickinson) and analyzed with FlowJo™ software; PI^+^ cells were excluded from analysis.

### Statistical analysis and normalization between experiments

For all bar graphs, dots represent values obtained from an individual experiment. Results between parasite strains were often expressed relative to the response elicited by the type II strain (equal 1), or the response to infected knockout macrophages normalized to infected wildtype macrophages (equal 1). All statistical analyses (one-way or two-way ANOVA with Bonferroni’s correction, and unpaired two-tailed t-test) were performed with GraphPad Prism version 8.3.0.

## ACKNOWLEDGEMENTS

We would like to thank Dominique Soldati-Favre (University of Geneva) for the ME49 *Δasp5* strain; Igor Brodsky (University of Pennsylvania) for *Gsdmd-/-* mouse bones; John Boothroyd (Stanford University) for anti-GRA7 polyclonal rabbit antibodies, ME49 *Δmyr1*, ME49 *Δmyr1::MYR1*, and RH *Δrop17* parasite strains; George Yap (Rutgers New Jersey Medical School) for sending the BOF +LC37 strain; Vishva Dixit (Genentech) for sending *Asc-/-* mice. We thank April Apostol (UC Merced) for initial help with IFA and visualization of TGD057-HA.

---

**Figure S1:**
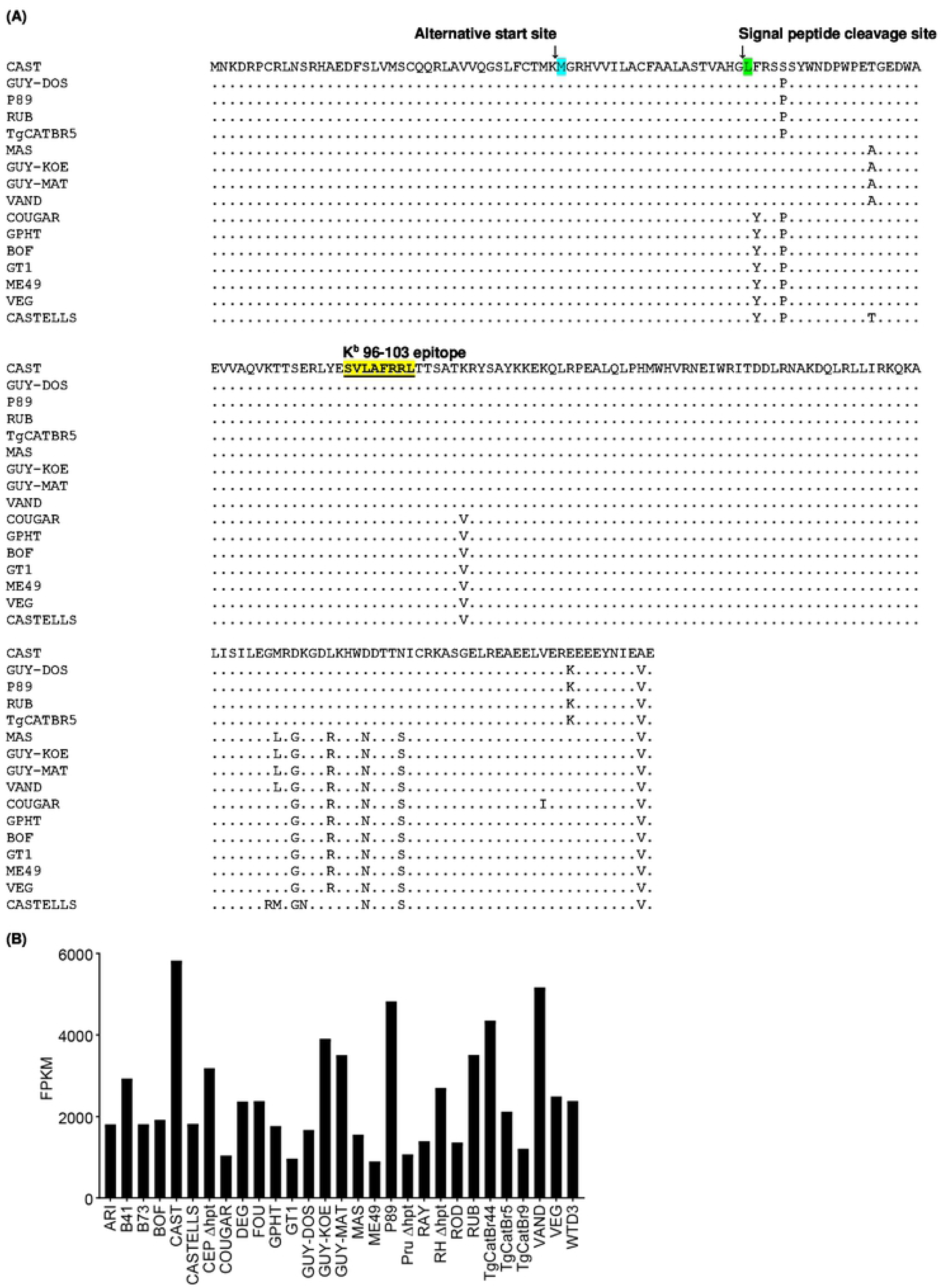
Conservation of the TGD057 96-103 peptide epitope and high *TGD057* gene expression between *T. gondii* strains. **(A)** Multiple protein alignment of TGD057 (215980) encoded by various *T. gondii* strains; the 96-103 MHC 1 K^b^ T57 T cell epitope is highlighted. Dots represent amino acid conservation with TGD057 from the CAST strain. The predicted signal peptide cleavage site and an alternative translational start site (54) is indicated with an arrow. **(B)** *TGD057* gene expression (Tg_215980) for 29 parasite strains following 20-22 hours post-infection in BMDMs (C57BL/6) is plotted from data previously reported (137); expression values are in fragments per kilobase of exon model, per million mapped reads (FPKM).

**Figure S2:**
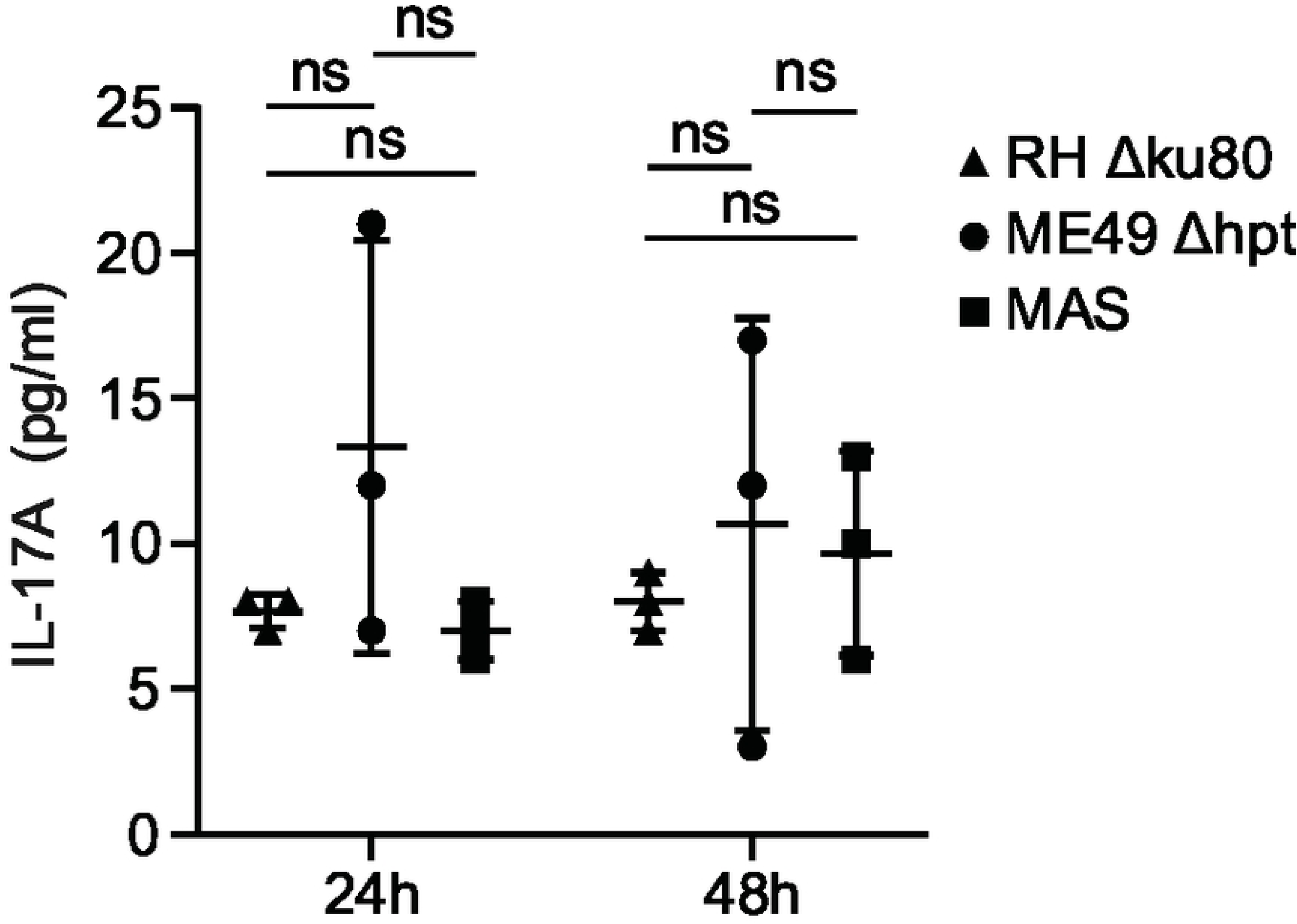
Negligible IL-17A production to *T. gondii*-infected BMDMs. BMDMs were infected with the indicated parasite strains and IL-17A was measured in the supernatant at 48h post addition of naïve T57 CD8 T cells. Average of 3 experiments + SD are plotted; each dot represents the result from an individual experiment. Statistical analysis was performed with two-way ANOVA with Bonferroni’s correction, ns non-significant.

**Figure S3:**
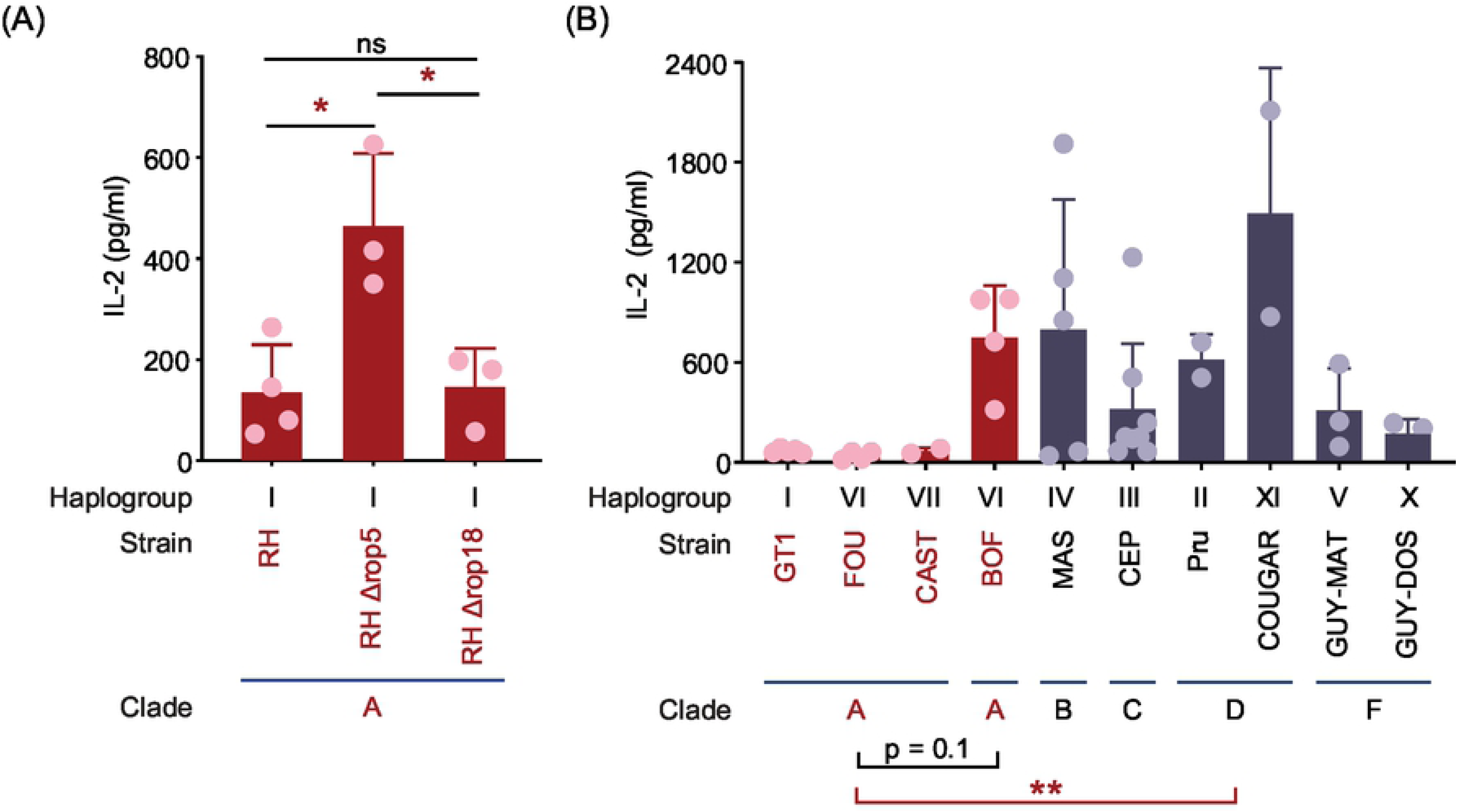
TGD057-specific CD8 T cell IL-2 responses to various strains of *T. gondii*. **(A)** BMDMs were infected with the indicated clade A RH *Δrop5* and *Δrop18* strains and IL-2 was measured in the supernatant at 48h post addition of naïve T57 CD8 T cells. Plotted is the average + SD of 3 experiments. Statistical analysis was performed using one-way ANOVA with Bonferroni’s correction; * p ≤ 0.05. **(B)** BMDMs were infected with *T. gondii* strains—clonal (types I-III), atypical (HG IV-X), and HG XI—representative of various clades and haplogroups. Infected BMDMs were incubated with naïve T57 CD8 T cells for 48 hours and IL-2 concentration in supernatant was measured by ELISA. Each dot represents the result from an individual experiment and the averages + SD of 2-8 experiments per strain are shown. Statistical analysis was performed using one-way ANOVA with Bonferroni’s correction comparing the grouped average of the indicated clade A strains against the grouped averages of clade B, C, D or F strains, or against the BOF strain; ** p ≤ 0.01.

**Figure S4:**
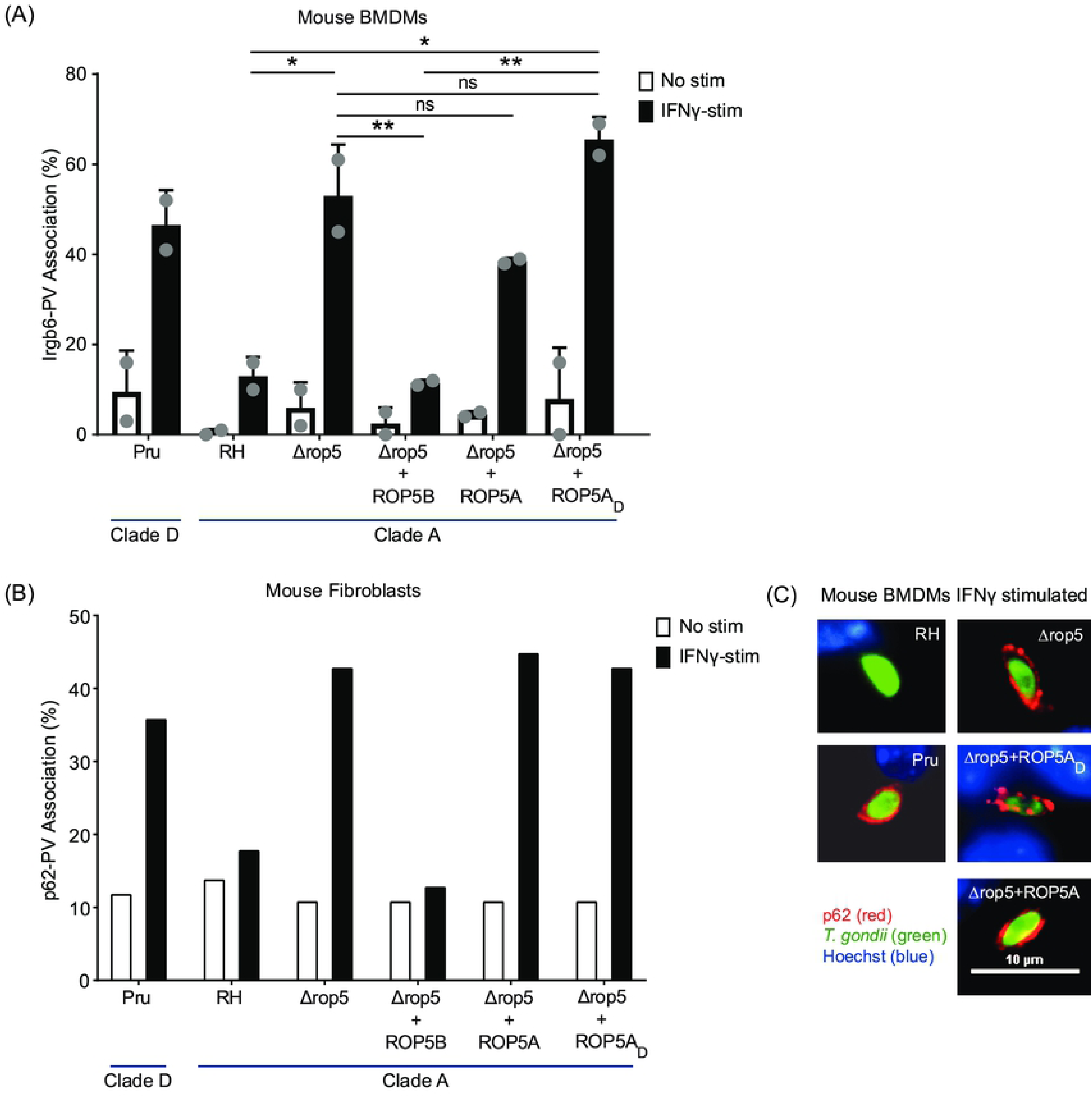
Irgb6- and p62-PV associations are not inhibited by *T. gondii* ROP5A isoforms. Untreated or IFNγ-stimulated BMDMs or MEFs were infected with indicated *T. gondii* strains. After 3-4 hours of infection, the samples were fixed and processed for IFA. *T. gondii* PVM was visualized in green with anti-GRA7 polyclonal rabbit antibodies. **(A)** Irgb6-PV localization was visualized in red with anti-Irgb6 polyclonal goat antibodies and plotted as percent of total GRA7+ *T. gondii* vacuoles. Plotted is the average + SD of 2 experiments. Statistical analysis was performed using one-way ANOVA with Bonferroni’s correction; * p ≤ 0.05, ** p ≤ 0.01, ns non-significant; shown are statistical comparisons between clade A strains. **(B)** P62-PV localization was visualized in red with a mouse anti-p62 monoclonal antibody and plotted as percent of total GRA7+ *T. gondii* vacuoles. **(C)** Representative images of p62-PV localization in parasite-infected IFNγ-stimulated BMDMs.

